# Plant interaction traits determine the biomass of arbuscular mycorrhizal fungi and bacteria in soil

**DOI:** 10.1101/2024.06.03.597074

**Authors:** Natascha Lewe, Robert A. Keyzers, Jason M. Tylianakis, Julie R. Deslippe

## Abstract

Plant-arbuscular mycorrhizal fungal (AMF) mutualisms play key roles in the biodiversity and productivity of ecosystems. Yet we have limited understanding of the functional roles of plants as AMF generalists or specialists, and the consequences of these plant interaction traits for soil ecosystems are virtually unknown. We grew eight pasture plant species under two experimental conditions and determined their root AMF communities by sequencing. We determined plant species interaction traits with AMF using a set of numeric and phylogenetic α-, β– and γ-diversities and characterized plant species’ relative interaction generalism for AMF. We used lipid analysis of rhizosphere soils and Bayesian modelling to explore how host interaction traits affected carbon allocation to AMF and bacteria. Plant interaction traits for AMF appear to be stable despite large variation in edaphic conditions and AMF pools. We show that host interaction generalism was associated with opposite patterns of bacterial and AMF biomass; the phylogenetic diversity of host interactions was positively associated with AMF biomass whereas the richness of host interactions was negatively associated with bacterial biomass in rhizosphere soils. Explicit consideration of plant interaction niches may enhance understanding of how changes in biodiversity affect ecosystem carbon cycling.

**Open research statement:** This publication does not use novel code, and the bioinformatic and statistic pipelines for this manuscript will be made available at https://github.com/NLewe/Bayes-interaction-niche.

Raw data will be made available at NCBI as BioProject: PRJNA997080

## Introduction

Species interactions are important for maintaining the biodiversity, productivity and resilience of ecosystems (Tylianakis et al. 2010; McCann 2000; Ratzke, Barrere, and Gore 2020). Many species depend on mutualistic relationships for crucial processes such as pollination, dispersal, resource acquisition or stress alleviation (Bascompte, Jordano, and Olesen 2006; Allesina and Tang 2012), such that these interactions comprise a key component of a species’ niche (Carscadden et al. 2020). A species’ interaction niche, the degree to which it interacts with members of another trophic guild, such as its mutualistic partners, is often described as a continuum between specialism and generalism, and has important ecological implications for both guilds (Poisot et al. 2015). For example, generalist pollinators tend to positively affects plant production (Maldonado, Lomáscolo, and Vázquez 2013), while specialists can enhance coexistence by reducing competition (Bastolla et al. 2009). In nature, the presence of a range of interaction niches contributes to biodiversity and community stability (Poisot et al. 2015; Dehling et al. 2021).

Despite this importance, the definition of a specialist and generalist is not always straightforward (Poisot et al. 2012; Rohr, Saavedra, and Bascompte 2014). For example, the term specialist is often applied to members of one guild that interact with few partners but also to those interacting with phylogenetically-related partners (Bascompte 2009; Montesinos-Navarro et al. 2015). By contrast, a generalist is commonly defined either as a species with many or diverse interactions. These differences in the definitions of specialism and generalism are problematic because they lead to either the pooling or the selection of a subset of species interaction traits, which may vary in their effects on the community.

Additionally, mutualistic interactions can be predicted through phylogenetic relationships (Rezende et al. 2007) or by species traits (Vázquez et al. 2009; Eklöf et al. 2013), and generality can be conserved across a species’ range (Emer et al. 2016), yet interactions (particularly those of generalists) can be determined by random encounter probability (related to species’ abundances; Vázquez et al. 2009) and shaped by the local environment (Tylianakis et al. 2008). Thus, it remains unclear to what extent the local environment shapes species interaction generalism. Resolving the interaction traits of mutualistic species would improve understanding of the assembly and maintenance of ecological communities.

Possibly the oldest mutualism among eukaryotes is that between plants and arbuscular mycorrhizal fungi (AMF), which occurs in more than three quarters of vascular plant species and most terrestrial ecosystems (Brundrett and Tedersoo 2018). Arbuscular mycorrhizal (AM) plants allocate 2-20% of photosynthetic carbon (C) (Konvalinková et al. 2017) to obligately biotrophic soil fungi of the subphylum Glomeromycotina (Spatafora et al. 2016). In exchange, plants receive multiple benefits from AMF, including improved water and nutrient acquisition (Vogelsang, Reynolds, and Bever 2006), and pathogen and stress resistance (Begum et al. 2019; Pu et al. 2022). Consequently, AM plants are significant sinks for atmospheric carbon dioxide (Parihar et al. 2020). While AMF abundance is highly correlated to soil C sequestration in field studies (Wilson et al. 2009), it is less clear how AMF diversity, largely mediated by plant hosts, influences carbon allocation to the soil microbial community. Enhanced understanding of plant interaction traits for AMF may provide insight to how host species affect the diversity and production of soil ecosystems (Bennett and Groten 2022).

Relative to other mutualisms, our understanding of plant interaction niches for AMF is in its infancy. This is due, in part, to several stochastic, abiotic and biotic filters on community assembly (HilleRisLambers et al. 2012) that are less easily disentangled in studies of AMF (Vályi et al. 2016). For example, root AMF communities vary according to the available AMF pool (Šmilauer et al. 2020), and plant species identity is key driver of variation in AMF composition and biomass (Veresoglou and Rillig 2014; Leff et al. 2018). Interestingly, host functional traits appear to correlate with their interaction niches. For example, in grasslands, grasses and forbs are associated with different AMF richnesses, colonisation rates and compositions, with grasses often hosting more taxa than forbs (Sepp et al. 2019; Šmilauer et al. 2020). Likewise, AMF colonisation rates covary with the major gradient of AM plant root traits (Bergmann et al. 2020). Hosts employ a range of strategies in selecting their AM mutualists because AMF taxa differ in their root colonisation patterns and nutrient transfer abilities (Lendenmann et al. 2011; Horsch, Antunes, and Kallenbach 2023). While generalist hosts may enjoy complementary effects of multiple AMF (Koide 2000; Jansa, Smith, and Smith 2008), trade-offs arise through increased C costs, especially if cheaters are present (Kiers and Denison 2008; Bever et al. 2009) and establishing specialist interactions with few beneficial AMF may be favourable in some environments (Werner and Kiers 2015).

However, plant hosts’ AMF community composition and diversity is context-dependent (Šmilauer et al. 2021; Tylianakis et al. 2008), which has impeded a general understanding on the relationships between plant interaction niches and production of soil ecosystems.

Distinct plant interaction niches for AMF have the potential to affect ecosystem C cycling directly by altering AMF communities and indirectly through AMF-mediated effects on soil bacterial communities. Up to 40% of photosynthetic C is lost from plant roots as fatty acids, carbohydrates and other metabolites, fuelling the growth of the AMF mycelium and a complex but specific community of rhizosphere bacteria (Marschner and Baumann 2003; Jiang et al. 2017). Arbuscular mycorrhizal fungi also produce metabolites that alter the bacterial composition of their hyphospheres (Huang et al. 2023) and nutrient availability in soil (Zhang et al. 2020). Collectively, these processes generate plant-soil feedbacks that alter subsequent plant community assembly processes (Crawford et al. 2019), determining ecosystem C cycling over larger spatial and temporal scales. So although plant interaction niches are important for determining the structure and function of soil communities, detailed knowledge of their effects on C allocation to AMF and bacterial communities remains sparse.

Here, we sought to characterise plant interaction niches with AMF and to learn how interaction traits affect AMF and bacterial biomass in rhizosphere soil. Firstly, we generated different biotic and abiotic filters on AMF community assembly by growing eight plant species under two experimental conditions. We test the hypothesis (**Hyp**_1_) that plant species’ interaction roles as AMF generalists or specialists are stable to these changes, comparing the multidimensional plant interaction niches under different experimental conditions by Procrustes analyses. Secondly, we sought to learn how plant interaction niches affect AMF biomass in rhizosphere soil. We expected generalist hosts to be capable of greater C allocation to AMF due to their enhanced nutrient supply resulting from complementarity effects of their AMF communities (Koide 2000; Jansa, Smith, and Smith 2008). In turn, we expected that higher rhizosphere AMF biomass would lead to greater root-encounter probability and a greater proportion of the root system being colonized, increasing interaction generalism. We therefore test the hypothesis (**Hyp**_2_) that host interaction generalism is positively associated with AMF biomass in rhizosphere soils. We quantified the abundance of the neutral lipid fatty acid (NLFA) 16:1ω5 as a proxy for AMF biomass, and modelled its response to plant interaction traits, accounting for plant phylogeny, roots and shoot biomass in a Bayesian framework. Finally, we explore the effect of host interaction generalism with AMF on bacterial biomass in the rhizosphere. While plant and AMF species may have differential effects on bacterial communities (Söderberg, Olsson, and Bååth 2002; Scheublin et al. 2010), we expected a positive relationship between soil bacterial biomass and plant interaction generalism due to complementary effects of many AMF on bacterial species. We therefore test the hypothesis (**Hyp**_3_) that soil bacterial biomass would increase in response to plants’ interaction traits associated with host generalism for AMF. We estimate bacterial biomass using phospholipid fatty acid (PLFA) analysis of bacterial biomarkers and model the effect of plant interaction traits, accounting for root and shoot biomass in a Bayesian framework. Our study reveals how plant interaction traits affect the productivity of soil ecosystems contributing to understanding of how changes in biodiversity affect ecosystem C cycling.

## Methods

### Study site and glasshouse experiments

We conducted two glasshouse experiments (which differed only in their soil and abiotic conditions) to characterise plant interaction niches for AMF and to test hypothesis 1, enabling us to determine whether interaction niches were sensitive to the soil and abiotic environmental conditions. We used experiment 2 to determine whether interaction traits affect AMF and bacterial biomass in rhizosphere soil. We selected eight co-occurring plant species from the pasture site where field soil was collected. In each experiment, five replicates per plant species were grown in mesocosms. Each mesocosm consisted of an individual plant seedling in a potting mix containing field-collected soil as source of AMF inoculum. We sought to generate different filters on AMF community assembly in experiments 1 and 2 by collecting field soil in different seasons and altering the potting mix composition. All mesocosms were maintained in a glasshouse for 16 weeks. At harvest, we collected the aboveground plant biomass and plant roots to determine their dry weight biomass, respectively. We sampled rhizosphere soil and randomly subsampled from the roots for later lipid extraction from both substrates and collected a small random subsample from the roots for later DNA extraction. For experiment 1, only root samples for DNA analysis were collected as described. Details on the study site, glasshouse experiment and harvest can be found in Appendix S1: Section S1.

### Characterising the AMF community

To identify AMF in plant root samples, we extracted DNA and amplified the internal transcribed spacer 2 (ITS2) region of the eukaryotic ribosomal DNA by polymerase chain reaction (PCR) using primers ITS3 and ITS4 (Tedersoo et al. 2014). The PCR products were sequenced on the Illumina MiSeq platform. The resulting amplicon sequence variants (ASVs) were assigned a fungal taxonomy using the UNITE 8.2 (2020) database (Nilsson et al. 2019). We filtered the data to contain only sequences assigned to the subphylum Glomeromycotina, analysing each ASV as a proxy for AMF species (Fu et al. 2022). See Appendix S1: Section S2 for details of molecular, bioinformatics and sampling completeness steps.

### Defining the plant-AMF interaction niche

Diversity of interactions partners has multiple components, and each can be measured differently (Morris et al. 2014). To comprehensively describe plant interaction niches for AMF, we calculated a suite of seven numeric and phylogenetic metrics of α-, β- and γ-diversity for AMF sequences derived from plant roots. These included mean AMF richness and Shannon’s diversity index per plant species (numeric α-diversities), and the average mean phylogenetic distance (MPD) for all replicates per species (phylogenetic α-diversity). We calculated the proportion of core AMF species (those present in ≥60 % of the replicates) per species (β(core), numeric β-diversity) and β(CU), (compositional units, phylogenetic β-diversity). Finally, pooling replicates per species, we calculated the total number of unique AMF (numeric γ-diversity) and total MPD (phylogenetic γ-diversity). Details of interaction niche metrics are in Appendix S1: Section S3.

### Stability of plant interaction niche for AMF

To examine if plant interaction niches for AMF were stable under different environmental conditions we compared plant-AMF interaction niches in experiments 1 and 2. For each plant species we created a table of experiment x absolute values of the diversity metrics and applied symmetric Procrustes analysis (Peres-Neto & Jackson, 2001) in *vegan* (Oksanen et al. 2022) to test the correlation of the plants’ interaction niches in the two experiments. A permutational test using the maximal number of permutations and the function *protest* was used to assess the statistical significance of each correlation. To visualise plant-interaction niches for AMF in the two experiments, diversity metrics were scaled to vary between 0 and 1, from most specialist to most generalist plant species and plotted as stacked radar plots.

### Quantifying soil microbial biomass

To test whether the plant interaction niche influenced C allocation to the soil microbial community, AMF and bacterial biomass were quantified using neutral lipid and phospholipid fatty acid (NLFA & PLFA) analysis. The fatty acid (FA) 16:1ω5 from the NL fraction correlates well with AMF structures in roots and soil (Sharma and Buyer 2015), and is a suitable proxy for C allocation to AMF by the host because AMF are unable to synthesize FAs but depend on the host for FA C14:0 (Luginbuehl et al. 2017). Bacterial biomass was estimated using 31 bacterial PLFA biomarkers (Appendix S1: Table S3). Lipids were extracted from lyophilized soil and root samples in experiment 2 following Lewe *et al*. (2021) with modifications described in Appendix S1: Section S4. AMF and bacterial biomass in soil and roots is reported as mean *±* standard deviation. Significant differences among groups were assessed using ANOVA or, when assumptions were not met, a Kruskal-Wallis test.

### Effect of host interaction niches on C allocation to AMF and bacteria

To test whether host interaction traits for AMF influenced C allocation to AMF and bacteria in rhizosphere soils, we used Bayesian linear mixed modelling. We applied principal component analysis (PCA) to extract uncorrelated linear recombinations of the diversity metrics used to characterise plant interaction niches, calculated per replicate. Principle component 1-3 collectively described 86.8% of the variation in plant niche space for AMF. We therefore modelled AMF and bacterial biomass in soils as a function of PC1, PC2 and PC3 of the PCA. To account for possible effects of plant biomass on C allocation to soil microbes, we included plant root and shoot biomass or their ratio (root:shoot) as co-variates in the models. Likewise, we included total AMF biomass of the plant’s root system as a covariate to account for possible differences in soil AMF biomass due to different root AMF biomass. Because the plant species were unequally grouped within three plant families, we accounted for phylogenetic non-independence among samples by including a phylogenetic covariance matrix as a random effect. We also included plant species identity as a random effect to account for plant functional traits unexplained by the phylogeny or root and shoot biomasses. The best fit model was selected based on convergence and accuracy criteria, including using posterior predictive checks and leave-one-out cross-validation (LOO-CV) from all possible variable configurations of the model (Vehtari, Gelman, and Gabry 2017). All Bayesian models were fitted in *brms* (Bürkner 2017). Details can be found in Appendix S1: Section S5.

## Results

### Plant species have stable interaction niches for AMF

The eight plant species differed in their interaction niches with AMF, as evidenced by large differences in the absolute (Appendix S2: Tables S3 – S5) and relative (Figure 1) values of diversity metrics among them. Also remarkable was the similarity of plant interaction niches when plants were grown under the two experimental conditions. Niches were similar despite that the plants hosted very different taxa and significantly different AMF communities in the two experiments (Appendix S2: Figure S3), suggesting considerable stability of interaction niche for AMF in the plant species we studied (Figure 1, Table 1). For example, in both experiments the grasses *H. lanatus* and *A. capillaris* were interaction generalists for AMF, with high values of numeric α-diversity, whereas members of the Asteraceae *A. millefolium* and *C. intybus* were specialists relative to the other plant species tested. In contrast, *S. arundinaceus* was a phylogenetic generalist characterized by high diversities, while the grass *P. cita* had intermediate interaction traits overall. Permutational Procrustes analysis of the absolute values of the numeric and phylogenetic *α-, β-* and *γ-*diversities for plant species confirmed that plant interaction niches with AMF were significantly correlated (except *C. intybus,* for which AMF detection was low) under the two experimental conditions (Table 1), suggesting stability of plant interaction niches for AMF, and supporting hypothesis 1.

**Figure 1:**
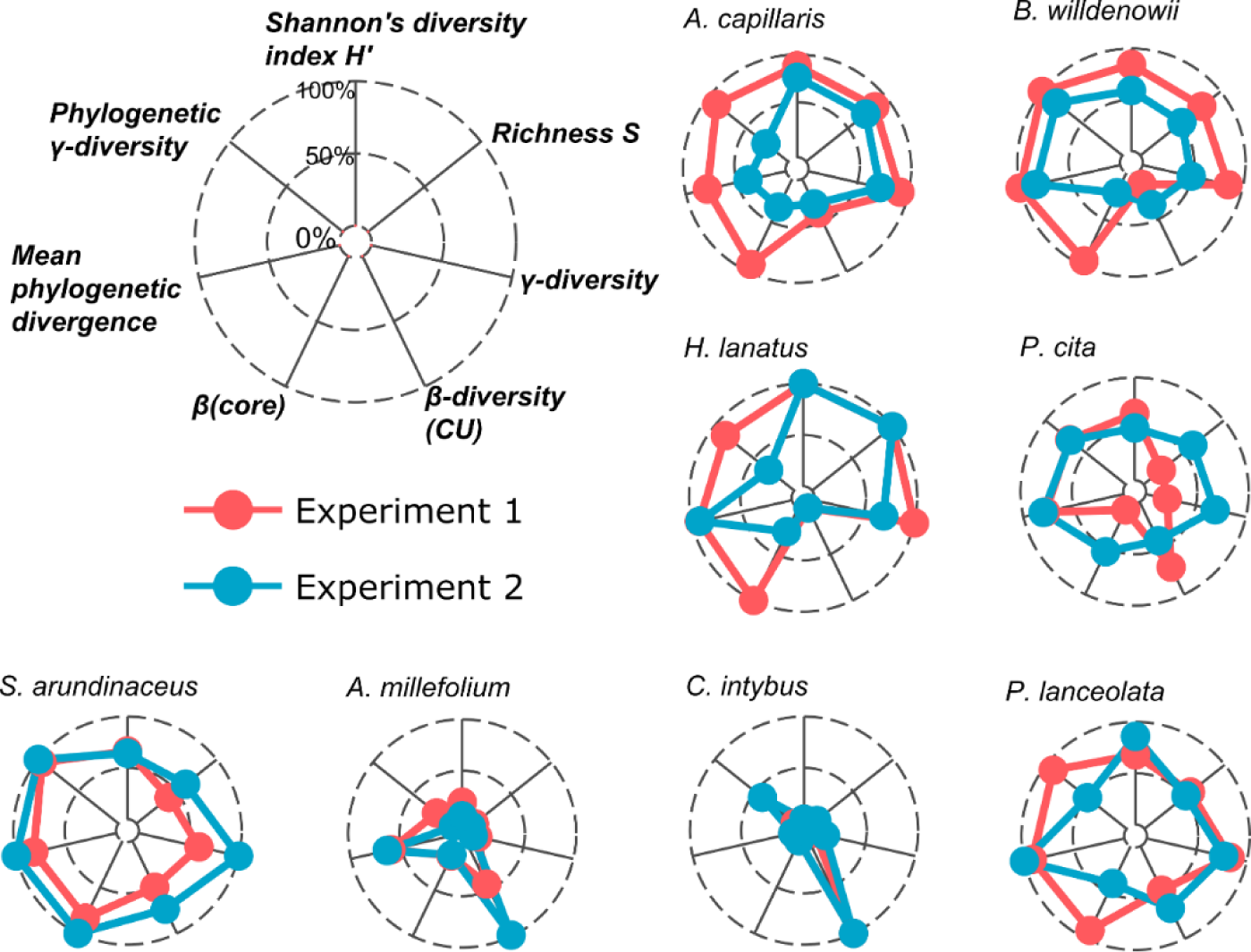
Radar plots comparing the relative interaction generalism of eight plant species as measured by seven metrics of numeric and phylogenetic α-, β- and γ-diversities under different experimental conditions (Experiment 1 and Experiment 2). We conceptualise the plant interaction niche for AMF as the area of the radar plot occupied by each plant species in each experiment. Values for each metric are the means per plant species (Appendix S2: Table S3) and were scaled between 0 and 1 for each experiment. CU = compositional units.

**Table 1:**
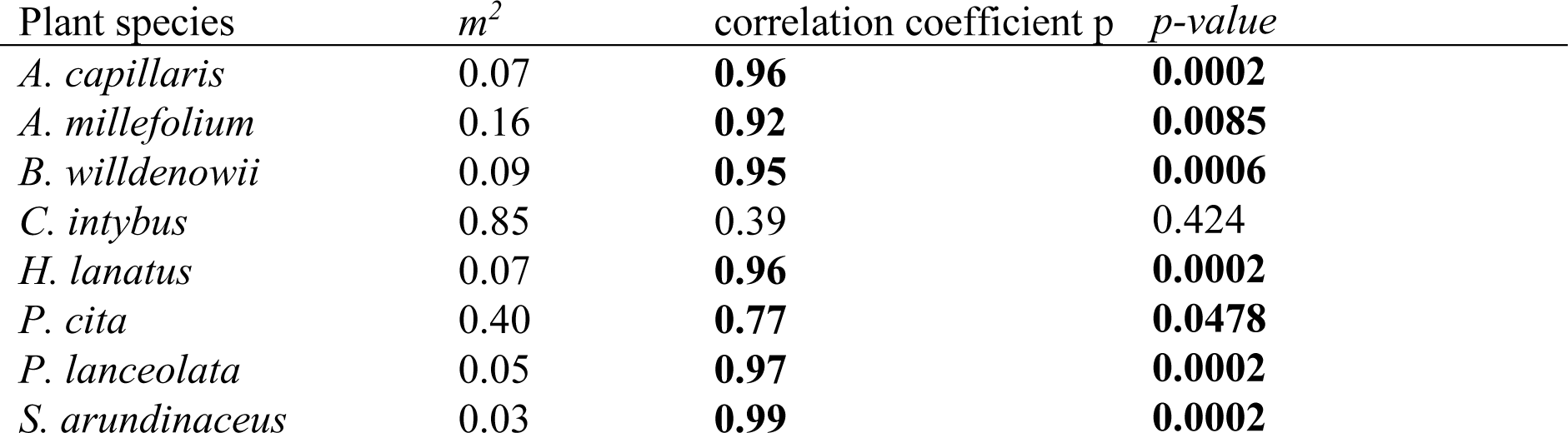
Permutational Procrustes analysis of plant interaction niches for AMF in experiment 1 and 2. Significant correlations are in bold. m^2^ = Procrustes sum of squares.

### Plant interaction generalism increases AMF biomass in rhizosphere soils

All plant species translocated large amounts of C into the AMF mycelium. The AMF biomarker NLFA 16:1ω5 was detected in all root and rhizosphere soil samples. Across rhizosphere soils, the AMF biomarker varied significantly from 5.7 ± 2.6 to 37.6 ± 13.1 nmol⋅g^-1^ DW soil (H = 15.8, df = 7, *p* = 0.027). In contrast, similar amounts of the AMF biomarker in the roots of all species suggests that they were similarly colonized by AMF (Appendix S2: Table S6) with values varying from 1.45 ± 0.62 μmol⋅g^-1^ DW root in *C. intybus* to 4.81 ± 1.39 μmol⋅g^-1^ DW root in *P. cita* (F (7, 28) = 2.35, *p* = 0.051).

The PCA of the diversity metrics used to characterize plant interaction niches for AMF revealed that numeric measures of α- and γ-diversity (richness, Shannon’s diversity, and γ-diversity) weighed heavily in forming PC1, which explained 52.6% of the variation in the plant-AMF interaction niche space. PC2 explained 22% of the variation and corresponded to phylogenetic γ-diversity (γ-MPD) and both metrics of β-diversity, β(CU) and β(core). PC3 explained an additional 12.2 % of the variance in plant-AMF niche space and was primarily composed of phylogenetic diversity (MPD) and β (CU) (see Appendix S2: Figure S4).

Our model for the AMF biomarker NL 16:1ω5 revealed that PC3 and plant root biomass had significant positive effects on AMF biomass in rhizosphere soil (Figure 2, Appendix S2: Section S4). The final model explained 34% of the variation of the AMF biomass in rhizosphere soils (Bayes R^2^ = 33.6 + 10.6 %). The retention of root biomass but not total AMF biomass of the root system as a co-variate in the final model indicated that C allocation to AMF is greatest for species with high biomass root systems regardless of their level of AMF colonization, while the positive effect of PC3 remains constant irrespective of root biomass. These results provide partial support to our hypothesis 2, indicating that while interaction-generalist hosts allocate more C to the soil AMF community, this relationship is strongly driven by phylogenetic interaction generalism and, to a much lesser degree, by β(CU).

**Figure 2:**
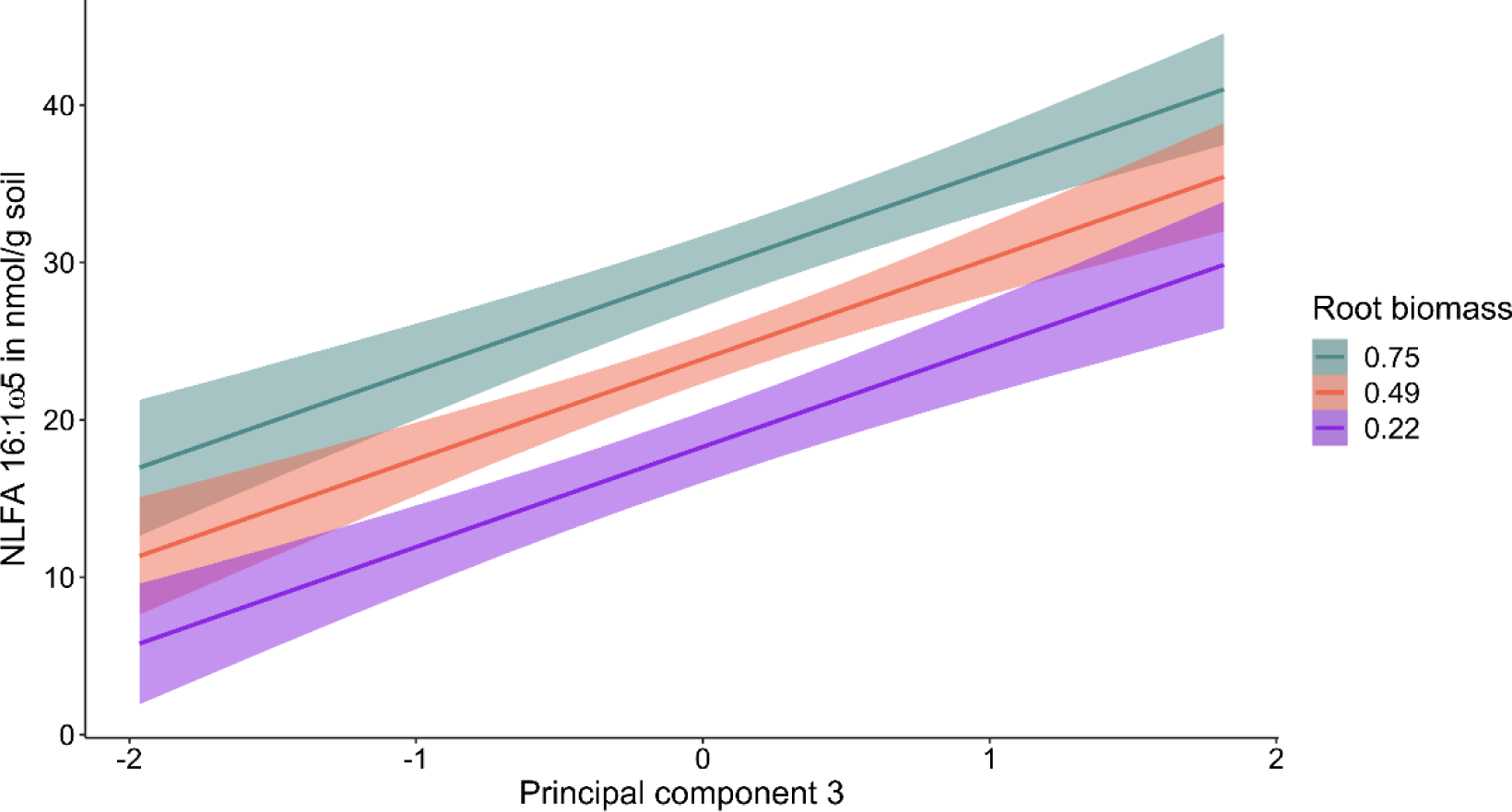
Predicted changes AMF biomass (NLFA 16:1ω5) in rhizosphere soil as a function of principal component 3 (PC3) and root biomass. PC3 represents phylogenetic y- and β-diversity, suggesting that these aspects of the plant interaction niche for AMF are associated with positive changes in AMF biomass in the rhizosphere. Root biomass was modelled as covariate, but here is visualised at three levels (mean root biomass ± standard deviation, uncertainty interval = 0.5). Final model: AMF biomass ∼ PC3 + root biomass.

### Plant interaction generalism for AMF reduces bacterial biomass in rhizosphere soil

Rhizosphere bacterial biomass varied significantly among plant species (H = 15.5, df = 7, p = 0.003) ranging from 14.66 ± 4.39 nmol⋅g^-1^ DW soil for *P. lanceolata* to 31.00 ± 6.30 nmol⋅g^-1^ DW soil for *C. intybus* (Appendix S2: Table S6). Contrary to our third hypothesis, our model for bacterial PLFAs revealed a strong negative effect of PC1 on bacterial biomass in rhizosphere soils, which was affected by plant shoot, and to a lesser degree, plant root biomass (Figure 3, Appendix S2: Section S5). The best model of soil bacterial biomass explained 49% of the variance (Bayes R^2^ = 49.1 ± 9.1%) and did not include a phylogenetic covariance structure or plant species identity as random effects.

**Figure 3:**
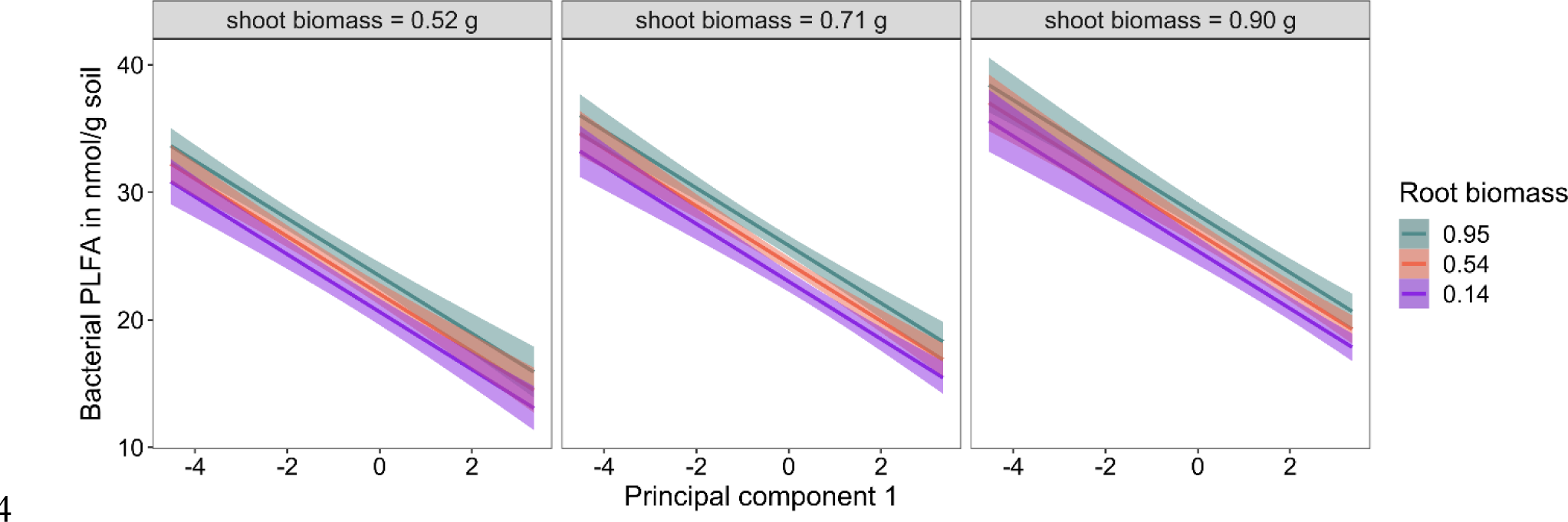
Predicted changes in soil bacterial biomass as a function of principal component 1 (PC1) modelled for three levels of shoot and root biomass, uncertainty interval = 0.5. PC1 represents numeric α- and γ-diversity (richness, Shannon’s diversity, and γ-diversity), suggesting that these aspects of the plant interaction niche for AMF are negatively associated with bacterial biomass in rhizosphere soils (Best model: soil bacterial biomass ∼ PC1 + root biomass + shoot biomass).

## Discussion

We found that pasture plant species exhibit fundamental interaction niches for AMF that are stable despite different environmental conditions. Our comprehensive characterisation of plant-AMF interaction niches provides novel insight to how plant niche partitioning for interaction partners affects C allocation to soil microbial communities. We show that C-allocation to AMF and bacteria were associated with different aspects of the plant interaction niche. Further, we show that interaction generalism had opposite effects on AMF and bacterial biomass in soils. Below, we discuss these results in detail and explore how plant interaction niches for AMF may affect ecosystem C cycling.

We found remarkable similarity in plant species’ interaction niches in experiment 1 and 2, despite that different edaphic conditions and AMF inoculum pool used in the two experiments generated substantial differences in the taxonomic composition of AMF communities of plant species, supporting our first hypothesis. Rhizosphere AMF communities change in response to edaphic conditions (Davison et al. 2021) and the available AMF pool (Van Geel et al. 2018), therefore our finding that plant interaction niches are relatively stable to these filters on community assembly suggests that plants exhibit fundamental interaction niches for AMF. Observations from plant-pollinator networks, where species retain their interaction niches moving from their native to alien ranges, corroborate these findings (Emer et al. 2016). However, given that we observed significant variation in the interaction niche of one of the eight species studied (*C. intybus*) and because we compared only two experiments, further work is needed to confirm the general stability of plant interaction traits for AMF. That said, given its direct influence on soil microbes, the characterisation of plant interaction generalism for AMF may be a useful functional trait (Funk et al. 2017) for understanding how the complex interplay of soil organisms affect ecosystem processes.

Our multidimensional approach to plant interaction generalism allowed us to resolve niche partitioning for AMF partners by different generalist plant hosts. We demonstrate further partitioning of interaction trait space by generalist hosts driven by variation in numeric and phylogenetic AMF diversities, which are likely to come with distinct trade-offs. Interaction niche partitioning may be affected through plant selection for the most beneficial AMF partners (Werner and Kiers 2015), or by interacting with AMF with a variety of nutrient acquisition and colonisation strategies (Powell and Rillig 2018). Plant niche partitioning for AMF partners may therefore contribute significantly to maintaining the functional diversity of ecosystems (Dehling et al. 2021), enhancing their resilience to environmental change (Turnbull et al. 2016).

We found some support for our second hypothesis that plant interaction generalism for AMF is positively related to AMF biomass in the rhizosphere. However, the increase in soil AMF biomass was driven by the phylogenetic diversity aspect of host interaction generalism. Phylogenetically diverse AMF communities are associated with higher trait variability like hyphal growth patterns (Hart and Reader 2002) and nutrient acquisition (Horsch, Antunes, and Kallenbach 2023). Therefore, our finding may suggest that complementarity among AMF taxa led to greater C allocation to the rhizosphere. Alternatively, interaction generalists hosting diverse AMF taxa may have been less able to downregulate C flow to less favourable mutualists (Grman 2012) making them more susceptible to cheaters (Kiers and Denison 2008). Indeed, the significant positive effect of β(CU), which reflects heterogeneity of AMF among replicates of a host species, on AMF biomass in the rhizosphere is congruent with the notion that generalist hosts may have been less able to select for beneficial AMF in our study. We found that root biomass was an important co-variate, whereas AMF biomass in the roots did not affect soil AMF biomass. While anatomical root traits, such as root diameter and branching are known to influence plant interaction niches for AMF (Bergmann *et al*., 2020; Ramana *et al*., 2023), our result may have been driven simply by more habitat for AMF in the hosts with higher root biomass (Sweeney *et al*., 2021). Higher root-habitat availability should reduce interspecific competition, supporting greater AMF diversity (Bergmann et al. 2020; Mony et al. 2021). Given the importance of AMF in C sequestration into the soil organic C pool (Zhu and Miller 2003), our result linking soil AMF biomass to the phylogenetic diversity aspect of plant interaction generalism highlights a key role for interaction-generalist plant species in regulating C flux between the atmosphere and biosphere.

Contrary to our third hypothesis, we found that interaction-generalist plants were associated with lower bacterial biomass in rhizosphere soils. The tri-partite interactions between plants, AMF and rhizosphere bacteria are complex, with both plants and AMF releasing compounds that can generate positive or negative effects on bacterial taxa (Bharadwaj, Alström, and Lundquist 2012; Changey et al. 2019). Furthermore, AMF and soil bacteria have overlapping niche space, and AMF can be effective competitors for resources in the rhizosphere. For example, the presence of AMF hyphae can significantly reduce bacteria competing for the same nutrients (Bukovská et al. 2018). Indeed, the effect of AMF on bacteria strongly depends upon the nutrient status of the host plant and AMF (Lanfranco, Fiorilli, and Gutjahr 2018; Huang et al. 2023). The positive effect of nutrient limitation on plant C-allocation to mycorrhizas is well known (Huang et al. 2023). It therefore seems plausible that under nutrient limitation, plants that host large AMF communities generate strong competitive effects on rhizosphere bacteria. In our study, plants and AMF were likely to be nutrient limited, as mesocosms were constructed of sand with only small volumes of field collected soil as inoculum and no mineral nutrient supplementation. Moreover, despite ample light, plants were relatively small at harvest, strongly suggesting nutrient limitation. Both root and shoot biomass were significant co-variates in our model of bacterial biomass, indicating that larger plants were associated with larger bacterial communities. Together, this suggests that competition for plant C among AMF and bacteria limited total bacterial biomass in our study.

We sought a greater understanding of plant interaction niches for AMF and how they may affect the biomass of soil microbial communities. We demonstrate that despite variation in mean values induced by edaphic contexts and the soil biota, plant species’ interaction niches for AMF were stable relative to other plants in their community, aligning with findings for plant functional traits generally (Funk et al. 2017) and for interaction traits in other types of networks (Emer et al. 2016). Under the nutrient-limited conditions of our experiments, we found that plants with high phylogenetic interaction generalism were associated with higher soil AMF biomass whereas high numeric interaction generalism drove lower bacterial biomass, suggesting strong AMF-bacterial competition for C in the rhizosphere. These findings are parsimonious with well-described patterns in community and ecosystem ecology, such as greater fungal: bacterial biomass (Wardle et al. 2004) and plant mycorrhizal dependence (Huang et al. 2023) under nutrient limitation. We suggest that plant interaction niches for AMF are a promising new avenue to enhance understanding of how plant traits alter key ecosystem functions, such as C cycling.

## Supporting information

Appendix2: Results

## Acknowledgements

This research was supported by a Marsden Fast Start Fund (15-VUW-069) awarded to JRD and JMT. NL was supported by a Victoria University of Wellington (VUW) Doctoral Scholarship. We appreciate Jan Vorster’s assistance with GC-MS inquiries, the team from ’Rāpoi,’ VUW’s High Performance Computing system, and Dr Lisa Woods for her valuable feedback on Bayesian modelling. We thank Maedeh Jafari Rad and the Deslippe lab group for their help in setting up and harvesting glasshouse experiment 2.

