## Appendix1: Methods for "Plant interaction traits determine the biomass of arbuscular mycorrhizal fungi and bacteria in soil"

### APPENDIX S1: ADDITIONAL METHODS

Lewe, Natascha, Deslippe, Julie R.. Plant interaction traits determine the biomass of arbuscular mycorrhizal fungi and bacteria in soil.

**Section S1:** Description of the study site, soil collection, glasshouse experiments and harvest

The study site from which we sourced soil for our experiments was a former pasture at Wairio Wetland (41°14’42’’S, 175°15’18’’E) on the Western shore of Lake Wairarapa, New Zealand. The collection area was previously used for pastoral agriculture but has been fenced off to exclude cattle in 2007 (Grant 2012). It represents a stable community consisting of mostly naturalised forb and graminoid species, with scattered stands of native kahikatea trees (*Dacrycarpus dacrydioides*) (Beadel et al. 2000, Gillon 2014). The soils of the study site are sandy and recent (Landcare Research 2016) with a silty texture in the upper 6 to 10 cm and a deep sandy layer (>1m) (Gillon 2014). The area is considered as relatively fertile (Sparling et al. 2008), with a total carbon content of 8%, and 0.75% nitrogen (N) (Landcare Research 2016). The inorganic phosphorus (“Olsen P”) falls between the low to moderate range with values between 10 and 25 µg mL$^{-1}$ (Sparling et al. 2008, Gillon 2014). The soil pH ranges from slightly acidic to circumneutral, varying between pH 6.0 to 7.2, while exhibiting a moderate cation-exchange capacity (25-40 cmolc kg$^{-1}$) (Gillon 2014, Landcare Research 2016).

Soil was collected at six randomly selected locations within the study site in spring (Experiment 2) and summer (experiment 1) in two different years. Field-collected soil was air-dried and sieved to remove plant roots and stones. This was to ensure the procurement of different AMF pools which would represent AMF pools the plants would encounter in the field. For experiment 1, the soil collection took place over the summer 2016/2017. Soil was homogenised by manual mixing on a tarpaulin in a 1:1 ratio with an autoclaved mix of peat, coarse sand and pumice (1:2:1, respectively). For experiment 2, collected in August 2018, the soil was mixed in a one to three ratio with a sand mix of coarse sand (Dalton’s washed sand No.2) and fine sand (Dalton’s washed sand No. 1) which had been mixed in a ratio of two to three.

**Table S1: Plant** species and seed origin used in experiment 1 and 2

| **Plant species** | **PlantFamily** | **Plant functional group** | **source of seeds** |
|---|---|---|---|
| *Achillea millefolium* | Asteraceae | Forbs | Kings Seeds No 4965 |
| *Agrostis capillaris* | Poaceae | Graminoids | PGG Wrightson ARR23TE |
| *Bromus willdenowii* | Poaceae | Graminoids | PGG Wrightson Cutivar: ATOM |
| *Cichorium intybus* | Asteraceae | Forbs | Kings seeds No. 4235 |
| *Holcus lanatus* | Poaceae | Graminoids | PGG Wrightson |
| *Plantago lanceolata* | Plantaginacea e | Forbs | PGG Wrightson TONOIAH Tonic plantain |
| *Poa cita* | Poaceae | Graminoids | nzseeds.co.nz |
| *Schedonorus arundinaceus* | Poaceae | Graminoids | Burnett's |

Seeds of eight pasture plants which co-occurred at our study site were sourced from agricultural suppliers (Table S1). The seeds were surface-sterilised in a 1% sodium hypochlorite solution, followed by 70% ethanol and distilled water for 10 min each before left to germinate on moistened filter paper in petri dishes in a temperature-controlled room (18°C with 12/12 hr day/night). Seeds were monitored regularly, and the filter paper moistened if needed. Petri dishes containing fungal contamination were discarded. Individual seedlings were transplanted to 2 L pots containing 1.7 L soil-mix when the first true roots appeared. Five replicate pots per plant species were established and grown in a glasshouse. Plants were watered automatically and the position of the pots on the benches randomised biweekly. Seedlings that developed from the seed bank were promptly removed. All plants were harvested after 4 months of growth. Roots and rhizosphere soil were kept frozen at -20°C and the shoots oven-dried for 2 days at 60°C until further processing.

**Section S2: Sample preparation, DNA extraction, PCR and sequencing**

After harvest, we thoroughly washed the root system of each individual plant in distilled water to remove soil. Using a scalpel, we cut roots into small segments (0.5 to 1 cm length) to subsample randomly for DNA extraction. Around 50 mg of pooled root segments were weighed and placed into a 1.5 mL Eppendorf tube containing steel beads (Green Eppendorf Lysis Kit, Next Advance, Inc., USA). The samples were homogenised for 2 x 30 s at 8,000 rpm (~ 6,000 X g) on a Precellys Evolution homogenizer (Bertin instruments, FR).

Next, we added 600 µL CTAB extraction buffer (2% hexadecyl(trimethyl)ammonium bromide, 1% polyvinylpyrrolidone (PVP40), 100 mM Tris(hydroxymethyl)aminomethane hydrochloride (tris-HCl), 1.4 M NaCl, 20 mM Ethylenediaminetetraacetic acid (EDTA)), and the samples incubated at 60°C for 1 hr (Doyle 1991). To ensure uniform treatment of the root homogenate we inverted the tube every 20 min. After incubation, we added 600 µL chloroform followed by centrifugation at 10,000 rpm (~ 9,200 X g) for 2 s (Eppendorf centrifuge 5415R, Germany). The aqueous layer was then transferred to a new 1.5 mL tube using a wide bore pipette tip. We carefully added 800 µL cold isopropanol and put the samples on ice for 10 min to precipitate the DNA. By centrifugation at 10,000 rpm for 1 min, the DNA was pelleted, and the supernatant was carefully discarded. We washed the DNA pellet twice with 800 µL of 80% ethanol by gently mixing, followed by centrifugation at 10,000 rpm (~ 9,200 X g) for 1 min. We discarded the supernatant and air-dried the pellet upside down at room temperature. The DNA pellet was then resuspended in 40 µL TE buffer (10 nM Tris-HCl, 1mM EDTA, adjusted to pH = 7.4). We assessed the quality and concentration of the DNA extracts by spectrophotometry (Nanodrop NP80, Implen, Germany) before freezing the DNA at -80°C.

To identify the AMF community, we amplified the internal transcribed spacer 2 (ITS2) region of the eukaryotic ribosomal DNA by polymerase chain reaction (PCR) using the modified versions of the primers ITS3 (5’-**CAHCGATGAAGAACGYRG**-3’) and ITS4 (5’-**TCCTSCGCTTATTGATATGC**-3’) (Tedersoo et al. 2014). Each primer included a 6-base barcode to enable multiplexing of the samples before sequencing (Table S2). Barcoded primers were ordered from Integrated DNA Technologies (IDT, Coralville, IO, USA. The 50 µL PCR reaction mix contained the extracted DNA at a concentration of 2 ng $\mu L^{-1}$, 25 µL of 2 x MyTaq$^{TM}$ Red Mix (Bioline, London, UK), 1 µL of each 10 µM primer and 8 µL 5 x TBT-Par to enhance the amplification (Samarakoon et al. 2013). Thermal cycling conditions were: 5 min at 95°C, then 35 cycles at 95°C for 45 s, 50°C for 45 s, and 70°C for 60 s; followed by 15 min at 72°C (GeneAmp PCR System 2700, Applied Biosciences, USA). We cleaned the PCR product with ExoSAP-IT™ (Thermo Fisher Scientific, USA) before sending it for sequencing on an Illumina MiSeq (Exp 1: AGRF, Melbourne, Australia, Exp 2: Macrogen, Australia) to generate paired-end reads of 300 bp. The image analysis for the generated data was performed in real time by the MiSeq Control Software (MCS) 2.6.2.1 and Real Time Analysis (RTA) 1.18.54. The Illumina bcl2fastq 2.20.0.422 pipeline was used to infer the sequence data.

***Table S2: Primer barcodes used for multiplexing of samples.*** *Combining the two barcoded forward primers with the 19 barcoded reverse primers resulted in 38 unique barcode combination, leading to the pooling of PCR products from 38 samples. The resulting pools were sent to a sequencing provider. Barcoded primers were synthesised by IDT (idtdna.com, IO, USA).*

| PRIMER | Barcode | Barcode sequence at 5’ end of primer | Primer code |
|---|---|---|---|
| Fungi Forward | barcode1 | ATCACG | ITS3_F_B1 |
| Fungi Forward | barcode3 | TTAGGC | ITS3_F_B2 |
| Fungi Reverse 1 | barcode25 | ACTGAT | ITS4_R_B3 |
| Fungi Reverse 2 | barcode4 | TGACCA | ITS4_R_B4 |
| Fungi Reverse 3 | barcode5 | ACAGTG | ITS4_R_B5 |
| Fungi Reverse 4 | barcode44 | TATAAT | ITS4_R_B6 |
| Fungi Reverse 5 | barcode7 | CAGATC | ITS4_R_B7 |
| Fungi Reverse 6 | barcode8 | ACTTGA | ITS4_R_B8 |
| Fungi Reverse 7 | barcode29 | CAACTA | ITS4_R_B9 |
| Fungi Reverse 8 | barcode35 | CATTTT | ITS4_R_B10 |
| Fungi Reverse 9 | barcode11 | GGCTAC | ITS4_R_B11 |
| Fungi Reverse 10 | barcode12 | CTTGTA | ITS4_R_B12 |
| Fungi Reverse 11 | barcode13 | AGTCAA | ITS4_R_B13 |

| | | | |
|---|---|---|---|
| Fungi Reverse 12 | barcode14 | AGTTCC | ITS4_R_B14 |
| Fungi Reverse 13 | barcode31 | CACGAT | ITS4_R_B15 |
| Fungi Reverse 14 | barcode16 | CCGTCC | ITS4_R_B16 |
| Fungi Reverse 15 | barcode17 | GTAGAG | ITS4_R_B17 |
| Fungi Reverse 16 | barcode18 | GTCCGC | ITS4_R_B18 |
| Fungi Reverse 17 | barcode19 | GTGAAA | ITS4_R_B19 |
| Fungi Reverse 18 | barcode45 | TCATTC | ITS4_R_B20 |
| Fungi Reverse 19 | barcode21 | GTTTCG | ITS4_R_B21 |

**Processing of raw sequence reads – bioinformatics**

Paired-end reads were demultiplexed and trimmed using *cutadapt* 3.2 (minimum primer overlap = 6; maximal error rate = 0.12) to remove the primer sequences. Cutadapt enables demultiplexing of paired-end reads in mixed orientation (*https://cutadapt.readthedocs.io/en/stable/guide.html#demultiplexing-paired-end-reads-in-mixed-orientation*) for which we choose a maximum overlap of 6 nucleotides and an error of 0. Demultiplexing was computed on the Rāpoi High Performance Compute Cluster (Victoria University of Wellington, NZ). We quality checked, merged and filtered the sequence reads using the DADA2 ITS pipeline 1.18 (*https://benjjneb.github.io/dada2/ITS_workflow.html*) in R 4.0.4 (R Core Team, 2021). For filtering, the DADA2 default parameters were applied except for the maximum allowance of errors for the reverse reads which we set at 4 (allowed error for forward reads = 2) due to their lower quality. We adjusted the error allowance as it allows for appropriate filtering of paired-end reads, because the truncation of reads to remove low quality regions is unsuitable for ITS sequences as they have variable lengths (Tedersoo et al. 2015). We included "pseudo-pooling" of the samples in the DADA2 method as it reduces the loss of sequence variants that might be rare in some samples but not in others. Inferring ASVs in pseudo-pooling mode includes an additional step in which each sample is compared against the whole set of ASVs of the data to identify rare reads. Next, we discarded chimeric reads using default parameters of the pipeline and we naïve Bayes trained the fungal ITS classifiers on the dynamic UNITE 8.2 (2020) reference database to assign the fungal taxonomy (Nilsson et al. 2019).

Next, we examined if the detected pool of fungal ASVs represents the total possible pool for each treatment by calculating inter- and extrapolated rarefaction and sampling completeness curves in R package *iNEXT* 2.0.20 (Chao et al. 2014, Hsieh et al. 2016). This extrapolation was based on estimated richnesses (Colwell et al. 2012) and included confidence intervals based on the bootstrap method (Chao & Jost, 2012. Additionally, we constructed a sampling coverage curve per plant species (and soil). Because measures of species richness tend to increase with sample completeness (Chao et al. 2014), we resampled the samples to the smallest common sample coverage (coverage-based rarefaction) in *metagMisc* 0.0.4 (Mikryukov 2022). All sampling curves were calculated in R package *iNEXT* (Chao et al. 2014, Hsieh et al. 2016), and we computed the standardisation to equal coverage using the function "phyloseq_coverage_raref" with 99 iterations in the package *metagMisc* 0.0.4 (*https://github.com/vmikk/metagMisc/*) on the Rāpoi High Performance Compute Cluster (Victoria University of Wellington, NZ). Lastly, we filtered the data to contain only sequences assigned to the subphylum Glomeromycotina, which are considered to be AMF. We defined each ASV that was identified as subphylum Glomeromycotina as being a proxy for an AMF species.

**Section S3: Description of the plant-AMF interaction niche**

*Table S2: Metrics used to describe the plants' interaction niche with AMF. For each interaction metric, a high value indicates an interaction generalist with AMF.*

| Metric | Formula | Description & interpretation |
|---|---|---|
| **Richness**<br><br>**S** | S = count of AMF species (i.e., ASVs) per plant individual. | Richness is the number of AMF the plant individual is interacting with. |
| **Shannon's diversity**<br><br>**H'** | H' estimates diversity, including the evenness of the AMF community per plant individual.<br><br>$H' = \sum_{i}^{n} p_i \ln(p_i)$<br><br>$p_i$ = proportional abundance of the AMF species $i$. | Increase in H' indicates an increase in either the number of AMF species or the evenness of their proportions among individuals per plant species, or both. |
| **γ-diversity** | $\gamma =$ count of total AMF species (i.e., ASVs) per plant species. | Describes the total AMF richness per plant species across all individuals. |

| Metric | Formula | Description & interpretation |
|---|---|---|
| **Compositional units**<br><br>**CU** | $CU = {}^{\gamma}/_{\bar{S}}$<br><br>$\gamma$ = total AMF richness per plant species<br>$\bar{S}$ = mean richness per plant species | Indicates the compositional heterogeneity of the samples, i.e., the variation in AMF composition between the individuals of each plant species (within-species diversity). |
| **Mean pairwise distance**<br><br>**MPD** | $MPD = \frac{\sum_{i}^{n}\sum_{j}^{n}\delta_{i,j}}{n}$<br><br>$i, j$ = AMF species<br>$i \neq j$<br>$n$ = number of branches of phylogenetic tree<br>$\delta$ = distance (i.e., phylogenetic branch length separating each AMF species) | MPD describes the phylogenetic dispersion, i.e., the phylogenetic distances among all species of the AMF community It is calculated per plant individual. |
| **Phylogenetic γ-diversity** | | The phylogenetic **γ**-diversity describes the phylogenetic dispersion of each plant species' AMF community. |
| **β(core)** | β(core) = $1 - {}^{\gamma_{60\,\%}}/_{\gamma}$<br><br>γ = total AMF species per plant species<br>$\gamma_{60\,\%}$ = total number of AMF species present in 60 % of all plant individuals. | A high value shows a low compositional similarity between the individuals of a plant species, indicative of a plant generalist. |

#### Numeric metrics - measures of α-diversity

We calculated the numeric interaction niche width of plants for arbuscular mycorrhizal fungi (AMF) by determining their richness (S) and Shannon's diversity index (H') using *phyloseq* 1.42.0 (McMurdie and Holmes 2013). To assess α-diversity, we utilized the arithmetic means per treatment as indicators. The richness (S) metric represents the numeric abundance, while Shannon's diversity incorporates the evenness of each species based on their relative abundances (Hill 1973). Lastly, we determined the total number of unique ASVs (γ-diversity) occurring across all replicates for each treatment.

#### Mean phylogenetic distance (MPD)

To investigate the phylogenetic diversities of the AMF communities as an indicator of phylogenetic interaction niche width, we aligned the sequences using *ClustalW* (Thompson et al. 1994) within the R package *msa* 1.30.0 (Bodenhofer et al. 2015). Subsequently, we calculated pairwise distances between the DNA sequences to construct a phylogenetic tree utilizing the neighbour joining algorithm, implemented in the R packages *seqinr* 4.2.236 (Charif and Lobry 2007) and *ape* 5.6.2 (Paradis and Schliep 2019).

The mean pairwise distance (MPD) characterizes the phylogenetic dispersion within a sample's AMF community by measuring the average branch length (Webb 2000) between each pair of AMF ASVs from a distance-based phylogenetic tree that includes all AMF species in the dataset. MPD was computed in *vegan 2.6-4* (Oksanen et al. 2022).

#### Compositional heterogeneity of the replicates - within treatment β-diversity

There are many different β-diversity measures that can describe the compositional heterogeneity of AMF sequence variants among treatment replicates (Tuomisto 2010, Anderson et al. 2011). In this study, we calculated the numeric β-diversity, specifically the number of compositional units (CU) per treatment ("true β-diversity"), using the formula $CU = \gamma / \overline{S}$ (Whittaker 1960, Tuomisto 2010).

To explore the variability among replicates while considering the identity of the interacting AMF species, we examined the proportional abundance of a treatment's core interacting species. β(core) was calculated by dividing the sum of core species per treatment, occurring in over 60% of their replicates, by the unique number of AMF species per treatment (γ-diversity).

**Section S4: Determination of the microbial biomass by phospholipid and neutral lipid fatty acid analyses (PLFA & NLFA)**

The identification of fatty acids (FAs) as biomarkers is a biochemical method that quantitatively delivers fatty acid profiles without the biases associated with culturing soil microbes or polymerase chain reaction (PCR) (Kirk et al. 2004).

Sample preparation

For FA analysis, all samples were lyophilised and a high-throughput phospho– and neutral lipid fatty acid (PLFA and NLFA) analysis was applied (Bligh and Dyer 1959, Buyer and Sasser 2012) as described in detail in Lewe *et al* (2021). In short, we extracted lipids from either about 100 mg root material or at least 600 mg soil with a one-phase chloroform:methanol:phosphate-buffer mixture spiked with the phospholipid 1,2-dinonadecanoyl-sn-glycero-3-phosphocholine (PL 19:0) and the neutral lipid 1,2,3- trinonadecanoyl-sn-glycerol (NL 19:0) as internal standards (each at 20 nmol per sample). The lipids were fractionated into neutral lipids (NLs), glycolipids and phospholipids (PLs) on a silica column. We derivatized the PLs and NLs by alkaline methanolysis to generate fatty acid methyl esters (FAMEs), which we concentrated under a nitrogen stream until dry and then dissolved in 75 µL hexane for further analysis on a Gas chromatography-mass spectrometry (GC-MS). The FAMEs were separated on a capillary column in a gas chromatograph equipped with a mass spectrometer (see Lewe et al., 2021 for parameters of GC-MS analysis).

Fatty acid methyl ester identification

For identification of the FAMEs, the mass spectra and retention times were compared with those of 31 commercially available standards (Matreya, USA; Nu-Chek, USA). We used the ω-reference convention X:YωZ for designation of the FAs (Frostegård and Bååth 1996); modified by Lewe et al. (2021, see Table S3). To determine the quantity of the analytes in response to the mass spectrometer used, the calculation of the response factor (RF) for each FAME is required (Dodds et al. 2005). Therefore, we obtained calibration curves for 25 FAME standards of the 31 selected FAMEs from two replicates each at five different concentrations and calculated the response factor (RF) for each FAME. We chose these FAME standards as they consisted of representatives from each structural group of interest (saturated, *iso*-branched, *anteiso*-branched, cyclic, monounsaturated, polyunsaturated). For FAMEs that could not be commercially obtained, the RF from the structurally most related FAME standard was applied.

Throughout the steps of the lipid analysis, lipid material is usually lost. To determine the amount of a FAME of interest in the original sample, we calculated the ratio of the FAME's RF to the RF of the internal standard, which resulted in the relative response factor of the FAME of interest. After FAMEs were identified and their peak area determined using the Shimadzu GC-MS solution software 4.44, we imported the data into R for all following calculations. Using the above equation, we calculated the amount of analyte per g dry weight (DW) sample, resulting in the molality b in mol·$g^{-1}$ DW substrate as measure of the amount per g DW soil. Although FAMEs are analysed with GC-MS, the substances of interests are FAs, derived either from extracted PLs or NLs as described before. Hereafter, we only use the terms PLFA or NLFA to describe these substances.

#### Biomarker Designation

A comprehensive set of 31 PLFA biomarkers was utilised, taking into consideration the large variability in the composition of microbial phospholipids (Table S3).

***Table S3: List of phospholipid fatty acid (PLFA) and neutral lipid fatty acid (NLFA) biomarkers utilised to estimate the bacterial and AM fungal biomass in the soil.*** *The nomenclature follows the omega-reference convention X:YωZ. X= number of the carbon atoms of the fatty acid, Y = number of double bonds, Z= position of the first double bond counted from the methyl end (omega-end) of the carbon chain. Prefixes: i= Iso branching, a= anteiso branching, cy = cyclopropane ring, 2OH = hydroxyl-group at position 2 counted from the alpha-carbon atom; 10Me = methyl group at position 10 counted from the a-carbon atom Suffixes: c= cis-configuration, t = trans-configurations.*

| **Microbial group** | **AMF** | **General bacteria** | **Gram-positive bacteria** | **Gram-negative bacteria** | **Actinobacteria** | **Fungi** |
|---|---|---|---|---|---|---|
| **Fatty acid biomarkers** | 16:1ω5c (PLFA) | 14:0 | i15:0 | 2OH10:0 | 10Me16:0 | 18:2w6 |
| | | 15:0 | a15:0 | 2OH12:0 | 10Me17:0 | 18:1ω9c |
| | 16:1ω5c (NLFA) | 16:0 | i16:0 | 3OH12:0 | 10Me18:0 | 18:3ω3 |
| | | 17:0 | a16:0 | 2OH14:0 | | |
| | | 18:0 | i17:0 | 3OH14:0 | | |
| | | | a17:0 | 2OH16:0 | | |
| | | | | 3OH16:0 | | |
| | | | | 16:1ω7c | | |
| | | | | 18:1ω7c | | |
| | | | | 18:1ω7t | | |
| | | | | 19:1ω9c | | |
| | | | | cy17:0 | | |
| | | | | cy19:0 | | |

References: (Bååth et al. 1992, Frostegård and Bååth 1996, Zelles 1997, Willers et al. 2015, Francisco et al. 2016)

#### Section S5: Bayesian modelling

##### Phylogenetic covariance structure for the plant species

To include information about the phylogeny of the studied species into model building, a phylogenetic tree of the eight plant species was fitted using a maximum likelihood approach. Tree building was done in R using the packages *ape* 5.6.2 (Paradis and Schliep 2019) and *phangorn* 2.10.0 (Schliep 2011) after alignment and concatenation of the gene sequences of the tRNA-Leu (trnl) gene and the matK gene (Table S4) in Jalview 2.11.2.6 (Waterhouse et al. 2009).The phylogenetic covariance was calculated using a model equal to Brownian motion (Felsenstein 1985). The calculated covariance structure was then used as “random effect” in all Bayesian linear mixed modelling approaches.

***Table S4: trnL and matK genes sourced from the NCBI database for building of the plants' phylogenetic tree***

| |
|---|
| Annotation for AY603268.1/1-835<br>*Achillea millefolium voucher Tribsch et al. 7823 (WU) trnL gene, partial sequence; trnL-trnF intergenic spacer, complete sequence; and tRNA-Phe (trnF) gene, partial sequence; chloroplast* |
| Annotation for EU119354.1/1-1471<br>*Agrostis capillaris trnT-trnL intergenic spacer, trnL gene, and trnL-trnF intergenic spacer, partial sequence; plastid; EMBL EU119354; MD5 aec6df6c4180eebe39c26331d358b9ad; RFAM RF00028* |
| Annotation for GU817987.1/1-928<br>*Cichorium intybus isolate 1155 tRNA-Leu (trnL) gene, partial sequence; trnL-trnF intergenic spacer, complete sequence; and tRNA-Phe (trnF) gene, partial sequence; plastid; EMBL GU817987; MD5 09e8b794e27bc04f6d3e0390256a4390; RFAM RF00028* |
| Annotation for AY101952.1/1-858<br>*Plantago lanceolata tRNA-Leu (trnL) gene, partial sequence; and trnL-trnF intergenic spacer region; chloroplast genes for chloroplast products* |
| Annotation for EF137606.1/1-983<br>*Holcus lanatus voucher Hodkinson25 TCD tRNA-Leu (trnL) gene, partial sequence; trnL-trnF intergenic spacer, complete sequence; and tRNA-Phe (trnF) gene, partial sequence; chloroplast; EMBL EF137606; MD5 5dc786bb97b8822b5752c15a6809b9b7; RFAM RF00028* |
| Annotation for GQ324407.2/758-1804<br>*Poa cita voucher OTA:Lloyd s.n. 058916 trnT-trnL intergenic spacer, partial sequence; tRNA-Leu (trnL) gene, complete sequence; and trnL-trnF intergenic spacer, partial sequence; chloroplast EMBL GQ324407, MD5 575f2b4b623f1f482ace63af2d09c2ba* |
| Annotation for EU395890.1/1-984<br>*Bromus catharticus voucher ICN* |
| Annotation for EU119373.1/419-1599<br>*Festuca arundinacea isolate P4078 trnT-trnL intergenic spacer, trnL gene, and trnL-trnF intergenic spacer, partial sequence; plastid; EMBL EU119373; MD5 21ba072def431370188db683b4dfd88f* |
| Annotation for OL537801.1/1-846<br>*Achillea millefolium voucher BRIT:Gostel406 maturase K (matK) gene, partial cds; chloroplast* |
| Annotation for KJ529327.1/1-969<br>*Agrostis capillaris voucher UZ 299.07 maturase K (matK) gene, partial cds; chloroplast* |
| Annotation for OL434897.1/1-788<br>*Bromus catharticus voucher GPAGA 269 maturase K (matK) gene, partial cds; chloroplast* |
| Annotation for MG225176.1/1-787<br>*Cichorium intybus voucher RBG-Blitz 004 maturase K (matK) gene, partial cds; chloroplast* |
| Annotation for HQ593296.1/1-793<br>*Festuca arundinacea voucher AP450 maturase K (matK) gene, partial cds; chloroplast* |
| Annotation for KJ529341.1/1-969<br>*Holcus lanatus voucher UZ 291.07 maturase K (matK) gene, partial cds; chloroplast* |
| Annotation for OL434917.1/1-776<br>*Plantago lanceolata voucher GPAGA 157 maturase K (matK) gene, partial cds; chloroplast* |
| Annotation for MW251101.1/1-788<br>*Poa cita voucher OTA:Lloyd sn OTA 058916 maturase K (matK) gene, partial cds; chloroplast* |

#### Principal component analysis (PCA)

The measures of diversity describing the plants' interaction niche and therefore their relative interaction generalism for AMF are strongly correlated. For removing this multicollinearity and extracting new variables describing the plants' relative interaction generalism, principal component analysis (PCA) was applied. For that, the numeric and phylogenetic metrics of α-, β- and γ-diversity per individual plant were included as variables for PCA which was calculated and visualized using the R packages *FactoMineR* 2.7 (Lê et al. 2008), *factoextra* 1.0.7 (Kassambara and Mundt 2020) and *tidyverse 2.0.0* (Wickham et al. 2019) using variables that were scaled to unit variance. The percentage contribution of each variable to each principal component was calculated and visualized. The top dimensions explaining 80% of the total variance were extracted for subsequent use as variables in modelling.

#### Fitting of Bayesian linear mixed models

Bayesian linear mixed models were fit and analysed using the R package *brms* 2.19.0 (Bürkner 2017) as interface for "Stan". For each model during the model selection process, 4 chains with 4,000 iterations, 1,000 as warm up and default (weakly informative) priors was run. Then, the models were compared by cross-validation using the leave-one-out criterium (LOO) and the function *loo_compare* from the *brms* R package (Vehtari et al. 2017). The models included the principal components PC1, PC2 and PC3, the DW root biomass and shoot DW biomass or the shoot:root ratio of the plants as fixed effects ("population-level factors") and the phylogenetic covariance structure and plant species identity as random factors (group level effects). All additive combinations of these factors were tested in models and the fixed factors were included in possible interactions as well. All models were tested by LOO cross-validation and the best model chosen provided it converged well. Convergence was assessed with the *loo* function and discarded if a Pareto k diagnostic value was estimated with $k > 0.7$ as this indicated impractical convergence rates. If several models showed a value for *loo* that did not significantly differ from the others (difference less than 4 (Sivula et al. 2022)), the model showing the best convergence criteria and predictive posterior checks was used. The quality of the convergence was also identified based on the number of divergent transitions encountered when running the iterations of the model. Divergent transitions can indicate that parameters are not accurately predicted.

All R code can be found in *https://github.com/NLewe/Bayes-interaction-niche*. The model fit was calculated using the *bayes_R2* function in package *brms*. The Bayes $R^2$, for Bayesian fits is calculated as variance of the predicted values divided by the variance of predicted values plus the expected variance of the errors (Gelman et al. 2019).
