## Appendix2: Results for "Plant interaction traits determine the biomass of arbuscular mycorrhizal fungi and bacteria in soil"

### Appendix 2: Supporting results

Lewe, Natascha, Keyzers, Robert A., Tylianakis, Jason M., Deslippe, Julie R.. Plant interaction traits determine the biomass of arbuscular mycorrhizal fungi and bacteria in soil.

#### Section 1: Sequencing results and rarefaction

***Table S1: Plant species and number of replicates used in experiment 1 and 2.*** *E1 R. = number of replicates obtained per treatment from glasshouse experiment 1. E2 R. = number of replicates obtained per treatment from glasshouse experiment 2. R. AMF = number of replicates in which Glomeromycotina could be detected after metabarcoding. AMF = Arbuscular mycorrhizal fungi.*

| **Plant species** | **Plant Family** | **Plant functional group** | **E1 R. (R. AMF)** | **E2 R. (R. AMF)** | **Source of seeds** |
|---|---|---|---|---|---|
| *Achillea millefolium* | Asteraceae | Forbs | 5 (4) | 5 (5) | Kings Seeds No 4965 |
| *Agrostis capillaris* | Poaceae | Graminoids | 4 (4) | 5 (5) | PGG Wrightson ARR23TE |
| *Bromus willdenowii* | Poaceae | Graminoids | 5 (5) | 5 (5) | PGG Wrightson Cutivar: ATOM |
| *Cichorium intybus* | Asteraceae | Forbs | 5 (3) | 5 (5) | Kings seeds No. 4235 |

| | | | | | |
|---|---|---|---|---|---|
| *Holcus lanatus* | Poaceae | Graminoids | 5 (5) | 5 (5) | PGG Wrightson |
| *Plantago lanceolata* | Plantaginaceae | Forbs | 5 (5) | 5 (5) | PGG Wrightson TONOIAH Tonic plantain |
| *Poa cita* | Poaceae | Graminoids | 3 (3) | 5 (5) | nzseeds.co.nz |
| *Schedonorus arundinaceus* | Poaceae | Graminoids | 5 (5) | 5 (5) | Burnett's |

***Figure S1: Rarefaction and sampling completeness curves of fungal amplicon sequence variant (ASV) richness for each plant species of experiment 1.*** *Solid line segments are calculated rarefaction and completeness curves, dotted line segments are extrapolations. Shaded areas in each plot show the 95% confidence interval of the extrapolation. B) Sample completeness curves by sequence reads. C) Sample coverage curves. Shaded areas in each plot show the 95% confidence interval of the extrapolation.*

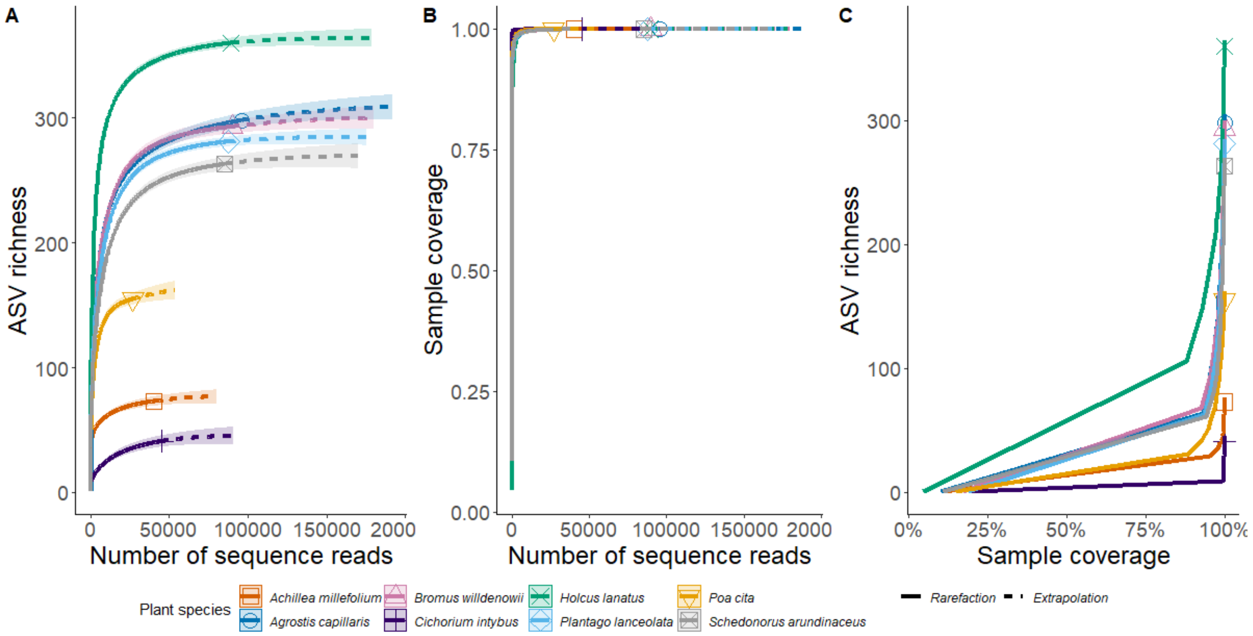

***Figure S2: Rarefaction and sampling completeness curves for fungal amplicon sequence variants (ASVs) by plant species, for roots for experiment 2.*** *Solid line segments are calculated rarefaction and completeness curves, dotted line segments are extrapolations. Shaded areas in each plot show the 95% confidence interval of the extrapolation. A), D) rarefaction sampling curves of fungal ASV richness. B), E) Sample completeness curves by sequence reads. C), F) Sample coverage curves in percent.*

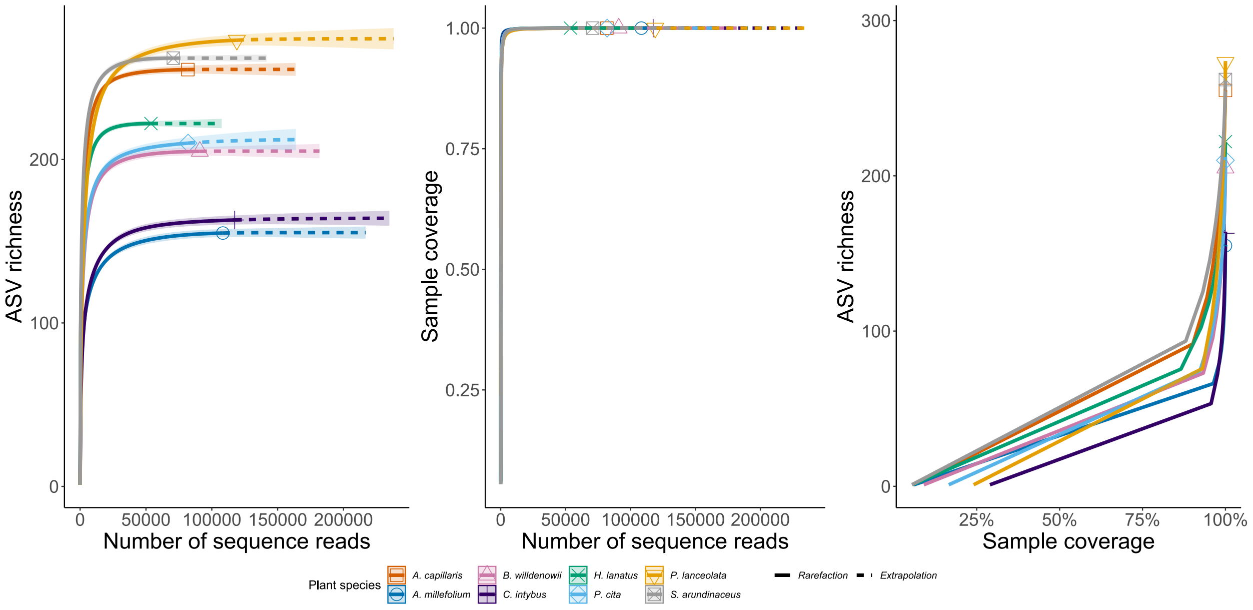

***Figure S3: Distribution of arbuscular mycorrhizal fungi (AMF) genera and absolute number of unique amplicon sequence variants (ASVs) per AMF genus, plant species and experiment (γ-diversity).*** *The experiments differed in their growth conditions, especially regarding their soil composition. Note the different number of replicates for A. capillaris (4) and P. cita (3) in experiment 1. E1 = Eperiment 1; E2 = Experiment 2.*

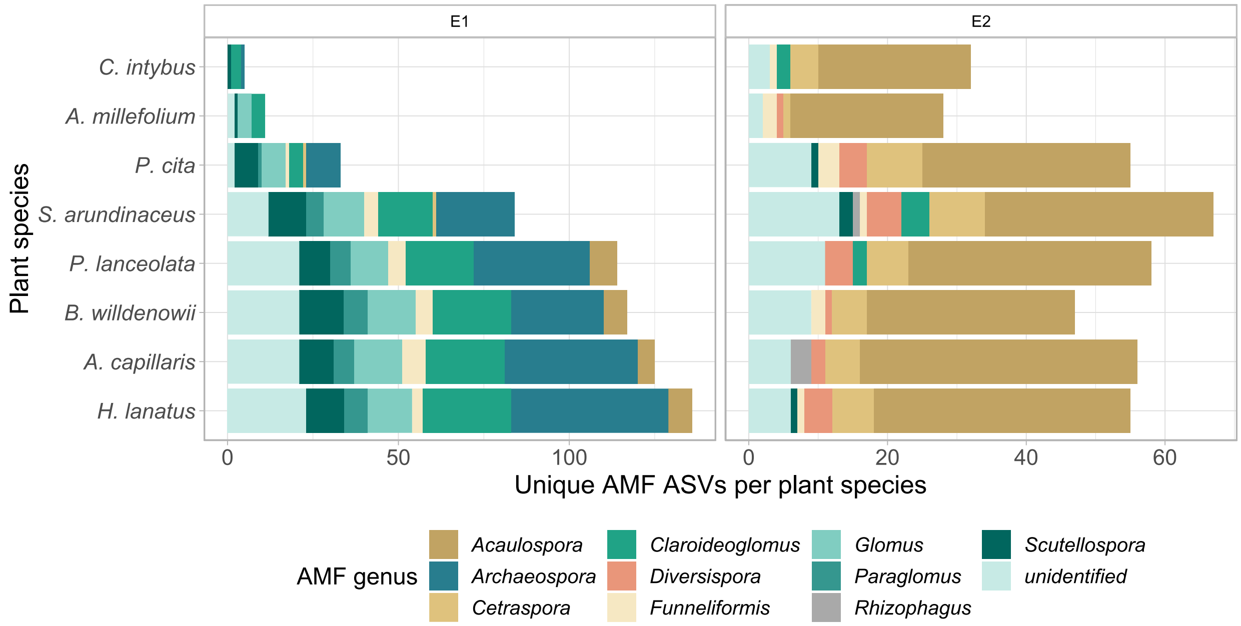

**Distribution of the Glomeromycotan families in experiment 1**

Overall, the frequencies of the ASVs and their distribution among the Glomeromycotan families varied greatly in experiment 1: 37% of the reads were assigned to the family Claroideoglomeraceae, followed by the Archaeosporaceae (26%), Acaulosporaceae (23%), Gigasporaceae (7%), Glomeraceae (4%), Paraglomeraceae (2%) and Diversisporaceae (0.04%). A total 0.02% of the read counts were unidentified at the family level. If considering the diversity of ASVs affiliated with these families, a different picture emerges. Here, the Archaeosporaceae dominate with a total of 55 detected sequences, followed by the Glomeraceae and Claroideoglomeraceae with 40 ASV each, the Acaulosporaceae (34 ASVs), Gigasporaceae (16 ASVs), Paraglomeraceae (9 ASVs), Diversisporaceae (1 ASV), and 1 sequence was unidentified at the family level.

**Distribution of the Glomeromycota in experiment 2**

The distribution of the Glomeromycotinan families varied greatly among the plant species' roots. In the roots, the Acaulosporaceae vastly dominated with 95% of the reads. The other identified families accounted for 2% (Gigasporaceae), 2% (Glomeraceae), 1% (Claroideoglomeraceae), 1% (Diversisporaceae), 0.05% (Archaeosporaceae) and 0.31% of the reads were unidentified at the family level. If considering the diversity of ASVs affiliated with these families, a different picture emerges with the Acaulosporaceae dominating only with a total of 77 of 124 detected sequences.

***Table S2: Number of sequence reads and amplicon sequence variants (ASVs) at different steps of the bioinformatics pipeline.*** *DADA = Divisive Amplicon Denoising Algorithm*.

| **Experiment 1 (root samples)** | **total** | **average per sample** |
|---|---|---|
| Initial sequence reads | 3,991,119 | 107,868 |
| Sequence reads after filtering and DADA2 pipeline | 1,528,330 | 41,306 |
| Number of ASVs after filtering and DADA2 pipeline | 733 | 90 |
| Sequence reads after rarefying to minimum coverage | 559,959 | 15,134 |
| Number of ASVs after removal of non-Glomeromycotinan ASVs | 196 | 34 |
| Number of sequence reads after removal of non-Glomeromycotinan ASVs | 19,103 | 562 |
| **Experiment 2 (root samples)** | | |
| Initial sequence reads | 1,178,093 | 31,002 |
| Sequence reads after filtering and DADA2 pipeline | 711,811 | 18,731 |
| Number of ASVs after filtering and DADA2 pipeline | 720 | 87 |
| Sequence reads after rarefying to minimum coverage | 158,848 | 3,971 |
| Number of ASVs after removal of non-Glomeromycotinan ASVs | 124 | 21 |
| Number of sequence reads after removal of non-Glomeromycotinan ASVs | 19,394 | 124 |

The dataset of experiment 1 included samples from 8 different plant species from five plant families (Table S1). For each plant species, five replicates were procured except for *A. capillaris* (four replicates), and *P. cita* (three replicates) for which the missing replicates did not reach harvesting age.

Consequently, sequencing results for 37 samples were obtained with an average of 107,868 reads per sample and a total read number of 3,991,119 (Table S2). After filtering, an average of 41,306 or 38% remained of the initial sequencing reads. The DADA2 pipeline resulted in a total number of 733 ASVs, and an average number of 90 ASVs per sample, ranging from 4 to 249 ASVs across all samples were assigned. As samples were rarefied to their minimal coverage, which was at 99.9% coverage, the average read count declined to 15,134 per sample. ASVs not belonging to the phylum Glomeromycota were removed from the data leaving 196 ASVs with a total count of 19,103 reads. This resulted in the removal of 3 samples (Appendix S1: Table S1), in which ASVs belonging to the Glomeromycotina could not be detected.

The dataset of experiment 2 consisted of 40 samples from roots of eight different plant species. Sequencing results of the fungal communities for those samples were obtained with an average of 31,002 reads per sample at a total read count of 1,178,093. 18,731 reads remained on average per sample after filtering. A total number of 711 ASVs resulted from the sample inference of the DADA2 pipeline with an average number of 87ASVs per sample. The rarefaction and sampling completeness curves (Appendix S2: Figure S2) showed that all plant species had been sufficiently sampled for fungal ASVs. After coverage-based rarefaction of the samples, a total of 158,848 reads with an average of 3,971 reads per sample remained. The removal of ASVs not belonging to the phylum Glomeromycota left 124 ASVs with a total count of 19,394 reads. Six samples were removed at that stage as they did not include any sequences after filtering for Glomeromycotina.

#### Section S2: Interaction niche metrics – detailed results

***Table S3: Values for all interaction metrics (α-, β- and γ-diversities) per plant species and experiment.*** *Values for the AMF richness S Shannon's H' and mean phylogenetic distance (MPD) are mean values per plant species, calculated by averaging the values over all plant individuals per species. Values for the β- and γ- diversities are defined on a per species level. The β-diversities are β(CU), i.e. the compositional units, and β(core) which describes the proportional abundance (between 0 and 1) of the core AMF (in 60% or more of the plant individuals, i.e. 3 of 5 replicates) interacting with the plant species. The γ- diversity describes the total number of AMF per plant species and the phylogenetic γ- diversity is calculated from uniting all replicates into one plant species sample and calculating the mean phylogenetic distance (MPD) for that plant species. E1 = experiment 1, E2 = experiment2.*

| Plant species | Shannon's H' | Richness S | γ-diversity | β(CU) | β(core) change 0 to 1 | Phylogenetic γ-diversity | UniFrac | MPD | Experiment |
|---|---|---|---|---|---|---|---|---|---|
| ***A. millefolium*** | 1.13 | 4.5 | 11 | 0.61 | 0.82 | 0.48 | 0.70 | 0.48 | E1 |
| ***A. capillaris*** | 3.15 | 53.5 | 125 | 0.58 | 0.79 | 0.55 | 0.62 | 0.51 | E1 |
| ***B. willdenowii*** | 3.01 | 48 | 117 | 0.49 | 0.72 | 0.56 | 0.54 | 0.54 | E1 |
| ***C. intybus*** | 0.58 | 2 | 5 | 0.83 | 0.8 | 0.46 | 0.77 | 0.37 | E1 |
| ***H. lanatus*** | 3.46 | 62 | 136 | 0.44 | 0.62 | 0.55 | 0.53 | 0.53 | E1 |
| ***P. lanceolata*** | 2.54 | 36.4 | 114 | 0.63 | 0.87 | 0.55 | 0.71 | 0.53 | E1 |
| ***P. cita*** | 2.45 | 15.33 | 33 | 0.72 | 0.67 | 0.53 | 0.62 | 0.51 | E1 |
| ***S. arundinaceus*** | 2.49 | 26.4 | 84 | 0.64 | 0.86 | 0.56 | 0.72 | 0.51 | E1 |
| ***A. millefolium*** | 1.72 | 8.8 | 28 | 0.64 | 0.86 | 0.37 | 0.59 | 0.33 | E2 |
| ***A. capillaris*** | 2.55 | 27.2 | 56 | 0.41 | 0.55 | 0.39 | 0.67 | 0.3 | E2 |
| ***B. willdenowii*** | 2.33 | 21.6 | 47 | 0.44 | 0.68 | 0.44 | 0.63 | 0.35 | E2 |

| | | | | | | | | | |
|---|---|---|---|---|---|---|---|---|---|
| ***C. intybus*** | 1.66 | 10 | 32 | 0.64 | 0.88 | 0.40 | 0.66 | 0.25 | E2 |
| ***H. lanatus*** | 2.8 | 33.6 | 55 | 0.33 | 0.33 | 0.40 | 0.42 | 0.36 | E2 |
| ***P. lanceolata*** | 2.64 | 21.6 | 58 | 0.54 | 0.76 | 0.41 | 0.58 | 0.37 | E2 |
| ***P. cita*** | 2.24 | 24 | 55 | 0.46 | 0.6 | 0.43 | 0.60 | 0.35 | E2 |
| ***S. arundinaceus*** | 2.39 | 24 | 67 | 0.56 | 0.79 | 0.46 | 0.61 | 0.37 | E2 |

***Table S4: Values for β(core) using a range of cutoffs.*** *β(core) was calculated by dividing the sum of core species per treatment, occurring in either one, two, three, four or all five of their five replicates (i.e., in ≥20%, ≥40%, ≥60%, ≥80% or a100% of the species' replicates), by the unique number of AMF species per treatment.*

| **Plant species** | **≥20% (1 of 5 samples)** | **≥40% (2 of 5 samples)** | **≥60% (3 of 5 samples)** | **≥80% (4 of 5 samples)** | **100% (5 of 5 samples)** | **Experiment** |
|---|---|---|---|---|---|---|
| *A. millefolium* | 0 | 0.55 | 0.82 | NA | NA | E1 |
| *A. capillaris* | 0 | 0.55 | 0.79 | 0.94 | 0.94 | E1 |
| *B. willdenowii* | 0 | 0.44 | 0.72 | 0.85 | 0.94 | E1 |
| *C. intybus* | 0 | 0.8 | 0.8 | NA | NA | E1 |
| *H. lanatus* | 0 | 0.39 | 0.62 | 0.76 | 0.95 | E1 |
| *P. lanceolata* | 0 | 0.6 | 0.87 | 0.95 | 0.99 | E1 |
| *P. cita* | 0 | 0.67 | 0.67 | 0.94 | 0.94 | E1 |
| *S. arundinaceus* | 0 | 0.63 | 0.86 | 0.94 | NA | E1 |
| *A. millefolium* | 0 | 0.68 | 0.86 | 0.93 | 0.96 | E2 |
| *A. capillaris* | 0 | 0.3 | 0.55 | 0.77 | 0.95 | E2 |
| *B. willdenowii* | 0 | 0.4 | 0.68 | 0.74 | 0.87 | E2 |
| *C. intybus* | 0 | 0.69 | 0.88 | 0.94 | 0.94 | E2 |
| *H. lanatus* | 0 | 0.16 | 0.33 | 0.62 | 0.84 | E2 |
| *P. lanceolata* | 0 | 0.47 | 0.76 | 0.93 | 0.98 | E2 |
| *P. cita* | 0 | 0.4 | 0.6 | 0.85 | 0.96 | E2 |
| *S. arundinaceus* | 0 | 0.51 | 0.79 | 0.94 | 0.97 | E2 |

***Table S5: Results for the numeric α-diversity metrics AMF richness S and Shannon's diversity H' per plant individual.*** *E1 = experiment 1, E2 = experiment.*

| **sample ID** | **Richness S** | **Shannon's H'** | **Plant species** | **Experiment** |
|---|---|---|---|---|
| AchMil1 | 2 | 0.693 | *A. millefolium* | E1 |
| AchMil2 | 1 | 0 | *A. millefolium* | E1 |
| AchMil4 | 8 | 1.96 | *A. millefolium* | E1 |
| AchMil5 | 7 | 1.875 | *A. millefolium* | E1 |
| AgrCap1 | 41 | 2.906 | *A. capillaris* | E1 |
| AgrCap2 | 68 | 3.694 | *A. capillaris* | E1 |
| AgrCap3 | 88 | 4.025 | *A. capillaris* | E1 |
| AgrCap5 | 17 | 1.981 | *A. capillaris* | E1 |
| BroWil1 | 51 | 2.86 | *B. willdenowii* | E1 |
| BroWil2 | 61 | 3.275 | *B. willdenowii* | E1 |
| BroWil3 | 27 | 2.706 | *B. willdenowii* | E1 |
| BroWil4 | 43 | 2.665 | *B. willdenowii* | E1 |
| BroWil5 | 58 | 3.568 | *B. willdenowii* | E1 |
| CicInt1 | 2 | 0.693 | *C. intybus* | E1 |
| CicInt2 | 3 | 1.04 | *C. intybus* | E1 |
| CicInt4 | 1 | 0 | *C. intybus* | E1 |
| HolLan1 | 49 | 2.859 | *H. lanatus* | E1 |
| HolLan2 | 62 | 3.547 | *H. lanatus* | E1 |
| HolLan3 | 107 | 4.165 | *H. lanatus* | E1 |
| HolLan4 | 40 | 3.177 | *H. lanatus* | E1 |
| HolLan5 | 52 | 3.568 | *H. lanatus* | E1 |
| PlaLan1 | 10 | 2.037 | *P. lanceolata* | E1 |
| PlaLan2 | 42 | 2.87 | *P. lanceolata* | E1 |
| PlaLan3 | 14 | 1.937 | *P. lanceolata* | E1 |
| PlaLan4 | 49 | 2.645 | *P. lanceolata* | E1 |
| PlaLan5 | 67 | 3.197 | *P. lanceolata* | E1 |
| PoaCit1 | 20 | 2.709 | *P. cita* | E1 |

| | | | | |
|---|---|---|---|---|
| PoaCit2 | 10 | 2.029 | *P. cita* | E1 |
| PoaCit3 | 16 | 2.612 | *P. cita* | E1 |
| SchAru1 | 30 | 2.981 | *S. arundinaceus* | E1 |
| SchAru2 | 19 | 2.588 | *S. arundinaceus* | E1 |
| SchAru3 | 5 | 1.461 | *S. arundinaceus* | E1 |
| SchAru4 | 53 | 3.281 | *S. arundinaceus* | E1 |
| SchAru5 | 25 | 2.157 | *S. arundinaceus* | E1 |
| R01 | 10 | 1.598 | *A. millefolium* | E2 |
| R02 | 8 | 1.455 | *A. millefolium* | E2 |
| R05 | 10 | 2.026 | *A. millefolium* | E2 |
| R09 | 7 | 1.822 | *A. millefolium* | E2 |
| R10 | 9 | 1.714 | *A. millefolium* | E2 |
| R12 | 19 | 2.415 | *C. intybus* | E2 |
| R13 | 2 | 0.693 | *C. intybus* | E2 |
| R14 | 15 | 2.131 | *C. intybus* | E2 |
| R15 | 11 | 2.011 | *C. intybus* | E2 |
| R18 | 3 | 1.028 | *C. intybus* | E2 |
| R21 | 29 | 1.949 | *P. cita* | E2 |
| R25 | 19 | 1.512 | *P. cita* | E2 |
| R26 | 31 | 2.84 | *P. cita* | E2 |
| R28 | 26 | 2.647 | *P. cita* | E2 |
| R30 | 15 | 2.247 | *P. cita* | E2 |
| R34 | 20 | 2.612 | *S. arundinaceus* | E2 |
| R36 | 34 | 2.298 | *S. arundinaceus* | E2 |
| R38 | 10 | 1.814 | *S. arundinaceus* | E2 |
| R39 | 35 | 3.167 | *S. arundinaceus* | E2 |
| R40 | 21 | 2.076 | *S. arundinaceus* | E2 |
| R42 | 36 | 2.992 | *H. lanatus* | E2 |
| R44 | 35 | 2.077 | *H. lanatus* | E2 |
| R45 | 36 | 3.061 | *H. lanatus* | E2 |
| R46 | 27 | 2.825 | *H. lanatus* | E2 |
| R47 | 34 | 3.064 | *H. lanatus* | E2 |

| | | | | |
|---|---|---|---|---|
| R51 | 37 | 2.755 | *A. capillaris* | E2 |
| R53 | 7 | 1.834 | *A. capillaris* | E2 |
| R54 | 32 | 2.693 | *A. capillaris* | E2 |
| R55 | 37 | 2.904 | *A. capillaris* | E2 |
| R59 | 23 | 2.563 | *A. capillaris* | E2 |
| R61 | 26 | 2.306 | *B. willdenowii* | E2 |
| R62 | 19 | 2.226 | *B. willdenowii* | E2 |
| R65 | 20 | 2.556 | *B. willdenowii* | E2 |
| R66 | 31 | 2.515 | *B. willdenowii* | E2 |
| R68 | 12 | 2.069 | *B. willdenowii* | E2 |
| R71 | 27 | 3.02 | *P. lanceolata* | E2 |
| R72 | 23 | 2.793 | *P. lanceolata* | E2 |
| R73 | 24 | 2.516 | *P. lanceolata* | E2 |
| R77 | 24 | 2.707 | *P. lanceolata* | E2 |
| R80 | 10 | 2.173 | *P. lanceolata* | E2 |

***Table S6: Mean phylogenetic distance (MPD) per plant individual.*** *Results significantly different from the null community are in bold. E1 = experiment 1, E2 = experiment 2.*

| sample ID | n (AMF) | MPD | mean MPD in null community | SD of MPD in null community | Standardised effect size of MPD vs. null communities | p-value of observed MPD vs. null communities | Experiment 1 or 2 |
|---|---|---|---|---|---|---|---|
| AchMil1 | 2 | 0.61 | 0.53 | 0.17 | 0.43 | 0.618 | E1 |
| AchMil2 | 1 | | | | | | E1 |
| **AchMil4** | **8** | **0.43** | **0.55** | **0.03** | **-3.82** | **0.004** | **E1** |
| **AchMil5** | **7** | **0.39** | **0.54** | **0.04** | **-4.09** | **0.002** | **E1** |
| AgrCap1 | 41 | 0.54 | 0.54 | 0.01 | -0.29 | 0.361 | E1 |
| **AgrCap2** | **69** | **0.53** | **0.54** | **0.01** | **-2.11** | **0.02** | **E1** |
| AgrCap3 | 88 | 0.55 | 0.55 | 0.01 | -0.58 | 0.265 | E1 |
| **AgrCap5** | **17** | **0.42** | **0.51** | **0.03** | **-3.29** | **0.005** | **E1** |
| BroWil1 | 51 | 0.55 | 0.55 | 0.01 | -0.13 | 0.442 | E1 |
| BroWil2 | 61 | 0.56 | 0.55 | 0.01 | 1.18 | 0.887 | E1 |
| BroWil3 | 27 | 0.54 | 0.54 | 0.01 | 0.07 | 0.506 | E1 |
| BroWil4 | 43 | 0.53 | 0.54 | 0.01 | -1.23 | 0.119 | E1 |
| BroWil5 | 58 | 0.54 | 0.55 | 0.01 | -1.17 | 0.13 | E1 |
| CicInt1 | 2 | 0.6 | 0.55 | 0.14 | 0.36 | 0.5395 | E1 |
| **CicInt2** | **3** | **0.15** | **0.52** | **0.11** | **-3.45** | **0.014** | **E1** |
| CicInt4 | 1 | | | | | | E1 |
| HolLan1 | 49 | 0.54 | 0.55 | 0.01 | -0.84 | 0.203 | E1 |
| HolLan2 | 62 | 0.53 | 0.54 | 0.01 | -2.09 | 0.022 | E1 |

| | | | | | | | |
|---|---|---|---|---|---|---|---|
| HolLan3 | 107 | 0.54 | 0.55 | 0 | -1.64 | 0.062 | E1 |
| HolLan4 | 40 | 0.55 | 0.55 | 0.01 | -0.07 | 0.468 | E1 |
| **HolLan5** | **52** | **0.5** | **0.54** | **0.01** | **-3.43** | **0.002** | **E1** |
| PhlPra4 | 16 | 0.53 | 0.53 | 0.02 | -0.14 | 0.434 | E1 |
| PhlPra5 | 18 | 0.56 | 0.55 | 0.02 | 0.58 | 0.692 | E1 |
| PlaLan1 | 10 | 0.56 | 0.55 | 0.03 | 0.48 | 0.651 | E1 |
| PlaLan2 | 42 | 0.54 | 0.54 | 0.01 | -0.31 | 0.365 | E1 |
| **PlaLan3** | **14** | **0.47** | **0.52** | **0.03** | **-2.21** | **0.02** | **E1** |
| PlaLan4 | 49 | 0.54 | 0.55 | 0.01 | -0.21 | 0.404 | E1 |
| **PlaLan5** | **67** | **0.53** | **0.54** | **0.01** | **-1.76** | **0.044** | **E1** |
| PoaCit1 | 20 | 0.54 | 0.54 | 0.02 | 0.05 | 0.485 | E1 |
| **PoaCit2** | **11** | **0.47** | **0.54** | **0.03** | **-2.35** | **0.025** | **E1** |
| PoaCit3 | 16 | 0.52 | 0.53 | 0.02 | -0.48 | 0.315 | E1 |
| SchAru1 | 30 | 0.57 | 0.55 | 0.01 | 1.46 | 0.946 | E1 |
| SchAru2 | 19 | 0.57 | 0.55 | 0.02 | 1.06 | 0.853 | E1 |
| **SchAru3** | **6** | **0.41** | **0.55** | **0.04** | **-3.39** | **0.008** | **E1** |
| SchAru4 | 53 | 0.53 | 0.54 | 0.01 | -1.44 | 0.078 | E1 |
| SchAru5 | 25 | 0.54 | 0.55 | 0.01 | -0.7 | 0.245 | E1 |
| R01 | 10 | 0.41 | 0.32 | 0.07 | 1.36 | 0.914 | E2 |
| R02 | 8 | 0.29 | 0.31 | 0.08 | -0.29 | 0.356 | E2 |
| R05 | 10 | 0.24 | 0.31 | 0.07 | -1.14 | 0.132 | E2 |
| R09 | 7 | 0.36 | 0.3 | 0.08 | 0.71 | 0.751 | E2 |
| R10 | 9 | 0.34 | 0.31 | 0.07 | 0.39 | 0.628 | E2 |
| R12 | 19 | 0.36 | 0.35 | 0.04 | 0.23 | 0.572 | E2 |
| R13 | 2 | 0.09 | 0.24 | 0.18 | -0.8 | 0.1555 | E2 |

| | | | | | | | |
|---|---|---|---|---|---|---|---|
| R14 | 15 | 0.4 | 0.33 | 0.05 | 1.25 | 0.888 | E2 |
| R15 | 11 | 0.3 | 0.32 | 0.06 | -0.29 | 0.368 | E2 |
| R18 | 3 | 0.11 | 0.27 | 0.14 | -1.08 | 0.1245 | E2 |
| R21 | 29 | 0.4 | 0.37 | 0.03 | 0.95 | 0.832 | E2 |
| R25 | 19 | 0.38 | 0.35 | 0.04 | 0.75 | 0.757 | E2 |
| R26 | 31 | 0.38 | 0.38 | 0.03 | 0.05 | 0.514 | E2 |
| **R28** | **26** | **0.29** | **0.36** | **0.03** | **-2.28** | **0.01** | **E2** |
| R30 | 15 | 0.28 | 0.33 | 0.05 | -1.1 | 0.145 | E2 |
| R34 | 20 | 0.31 | 0.35 | 0.04 | -1 | 0.173 | E2 |
| **R36** | **34** | **0.49** | **0.39** | **0.03** | **4.06** | **1** | **E2** |
| R38 | 10 | 0.3 | 0.32 | 0.07 | -0.24 | 0.412 | E2 |
| R39 | 35 | 0.43 | 0.39 | 0.02 | 1.83 | 0.974 | E2 |
| R40 | 21 | 0.32 | 0.35 | 0.04 | -0.78 | 0.214 | E2 |
| R42 | 36 | 0.36 | 0.39 | 0.02 | -1.16 | 0.128 | E2 |
| R44 | 35 | 0.4 | 0.39 | 0.02 | 0.62 | 0.733 | E2 |
| **R45** | **36** | **0.31** | **0.39** | **0.02** | **-3.38** | **0.001** | **E2** |
| R46 | 27 | 0.41 | 0.37 | 0.03 | 1.39 | 0.917 | E2 |
| **R47** | **34** | **0.32** | **0.38** | **0.02** | **-2.77** | **0.002** | **E2** |
| R51 | 37 | 0.37 | 0.39 | 0.02 | -0.91 | 0.188 | E2 |
| R53 | 7 | 0.2 | 0.3 | 0.08 | -1.23 | 0.102 | E2 |
| **R54** | **32** | **0.33** | **0.38** | **0.03** | **-1.88** | **0.032** | **E2** |
| **R55** | **37** | **0.32** | **0.39** | **0.02** | **-3.07** | **0.001** | **E2** |
| **R59** | **23** | **0.26** | **0.36** | **0.04** | **-2.75** | **0.008** | **E2** |
| R61 | 26 | 0.39 | 0.37 | 0.03 | 0.84 | 0.785 | E2 |
| R62 | 19 | 0.31 | 0.35 | 0.04 | -0.89 | 0.195 | E2 |

| | | | | | | | |
|---|---|---|---|---|---|---|---|
| **R65** | **20** | **0.28** | **0.35** | **0.04** | **-1.75** | **0.038** | **E2** |
| R66 | 31 | 0.34 | 0.38 | 0.03 | -1.4 | 0.087 | E2 |
| **R68** | **12** | **0.44** | **0.32** | **0.06** | **1.95** | **0.984** | **E2** |
| R71 | 27 | 0.35 | 0.37 | 0.03 | -0.41 | 0.331 | E2 |
| **R72** | **23** | **0.44** | **0.36** | **0.04** | **2.35** | **0.988** | **E2** |
| R73 | 24 | 0.37 | 0.36 | 0.04 | 0.4 | 0.639 | E2 |
| R77 | 24 | 0.33 | 0.36 | 0.04 | -0.69 | 0.246 | E2 |
| R80 | 10 | 0.35 | 0.32 | 0.07 | 0.44 | 0.661 | E2 |

#### Section S3: Biomass estimates

***Table S7:*** *Soil and root AMF biomass estimates, bacterial biomass estimates, shoot and root biomass. AMF and bacterial biomass estimated by NLFA and PLFA per g dry weight substrate (either roots or soil). Root and shoot biomass are in g dry weight. NA = sample lost during lipid extraction.*

| **sample ID** | **AMF biomass estimate (NLFA 16:1ω5 in nmol per g soil)** | ***AMF biomass estimate (*NLFA 16:1ω5 in µmol per g root)** | ***Plant species*** | **total bacterial biomass estimate (total bacterial PLFA biomarker in nmol per g soil)** | **shoot DW in g** | **root DW in g** |
|---|---|---|---|---|---|---|
| R01 | 32.5 | *2.26* | *A. millefolium* | 26.52 | 0.417 | 0.272 |
| R02 | 35.45 | *5.77* | *A. millefolium* | 32.75 | 0.5 | 0.234 |
| R05 | 27.22 | *3.45* | *A. millefolium* | 29.77 | 0.435 | 0.396 |
| R10 | NA | *2.26* | *A. millefolium* | NA | 0.549 | 0.407 |
| R12 | 40.3 | *2.24* | *C. intybus* | 29.51 | 0.6 | 0.719 |
| R13 | 18.87 | *1.06* | *C. intybus* | 27.82 | 0.385 | 0.612 |
| R14 | 34.84 | *0.88* | *C. intybus* | 31.29 | 0.529 | 1.047 |
| R15 | 55.4 | *2.00* | *C. intybus* | 24.92 | 0.71 | 1.335 |
| R18 | 38.68 | *1.07* | *C. intybus* | 41.47 | 0.552 | 2.237 |
| R21 | 5.97 | *5.70* | *P. cita* | 28.8 | 0.841 | 0.129 |
| R25 | 2.54 | *6.00* | *P. cita* | 27.36 | 0.799 | 0.328 |
| R26 | 8.8 | *2.92* | *P. cita* | 14.75 | 0.541 | 0.267 |
| R30 | 5.53 | *4.61* | *P. cita* | 23.3 | 0.842 | 0.092 |
| R34 | 10.01 | *3.95* | *S. arundinaceus* | 16.28 | 0.78 | 0.621 |
| R36 | 67.96 | *2.54* | *S. arundinaceus* | 17.8 | 0.641 | 0.539 |
| R39 | 48.77 | *0.86* | *S. arundinaceus* | 25.75 | 1.111 | 0.661 |

| | | | | | | |
|---|---|---|---|---|---|---|
| R40 | 6.29 | *2.44* | *S. arundinaceus* | 27.21 | 0.705 | 0.702 |
| R42 | 26.57 | *2.53* | *H. lanatus* | 28.31 | 0.764 | 0.453 |
| R44 | 35.98 | *3.05* | *H. lanatus* | 27.66 | 1.048 | 0.409 |
| R45 | 29.91 | *2.16* | *H. lanatus* | 25.2 | 0.636 | 0.413 |
| R46 | 10.97 | *0.47* | *H. lanatus* | 14.21 | 0.688 | 0.435 |
| R47 | 30.93 | *1.71* | *H. lanatus* | 22.45 | 0.996 | 0.6 |
| R51 | 31.08 | *2.15* | *A. capillaris* | 21.49 | 0.657 | 0.603 |
| R53 | 19.92 | *4.71* | *A. capillaris* | 22.83 | 0.801 | 0.23 |
| R54 | 20.1 | *2.16* | *A. capillaris* | 20.88 | 0.907 | 0.585 |
| R55 | 9.91 | *2.11* | *A. capillaris* | 25.25 | 0.803 | 0.348 |
| R59 | 13.33 | *0.29* | *A. capillaris* | 24.32 | 0.634 | 0.809 |
| R61 | 8.28 | *2.61* | *B. willdenowii* | 25.13 | 0.855 | 0.239 |
| R62 | NA | *2.52* | *B. willdenowii* | 14.94 | 0.737 | 0.249 |
| R65 | 25.73 | *1.64* | *B. willdenowii* | 20.01 | 0.7 | 0.21 |
| R66 | 23.1 | *1.75* | *B. willdenowii* | 20.66 | 1.026 | 0.445 |
| R68 | 23.52 | *2.83* | *B. willdenowii* | 23.13 | 0.805 | 0.305 |
| R71 | 13.04 | *3.34* | *P. lanceolata* | 9.74 | 0.412 | 0.418 |
| R72 | 31.77 | *8.11* | *P. lanceolata* | NA | 0.55 | 0.456 |
| R73 | 15.4 | *2.67* | *P. lanceolata* | 18.19 | 0.734 | 0.683 |
| R80 | NA | *1.63* | *P. lanceolata* | NA | 0.373 | 0.484 |
| Sc1 | 0.0 | NA | *Soil control* | 10.3 | NA | NA |
| Sc2 | 0.39 | NA | *Soil control* | 19.0 | NA | NA |
| Sc3 | 0.56 | NA | *Soil control* | 12.8 | NA | NA |

#### Section S4: PCA

***Figure S4: Principal component analysis (PCA) of numeric and phylogenetic α-, β- and γ-diversities per plant individual.*** *A & B) PCA-biplots. Dots are plant individuals, colored by plant species; blue arrows are metrics of diversity. A) PCA for PC1 and PC2. B) PCA for PC2 and 3. C)-E) Contributions of metrics to the first three principal components (PCs). The red dashed line corresponds to the expected value if the metrics had uniform contributions. β(core) = percentage of AMF species occurring in 60% of replicates per plant species, MPD = mean phylogenetic distance, β(CU) = compositional units.*

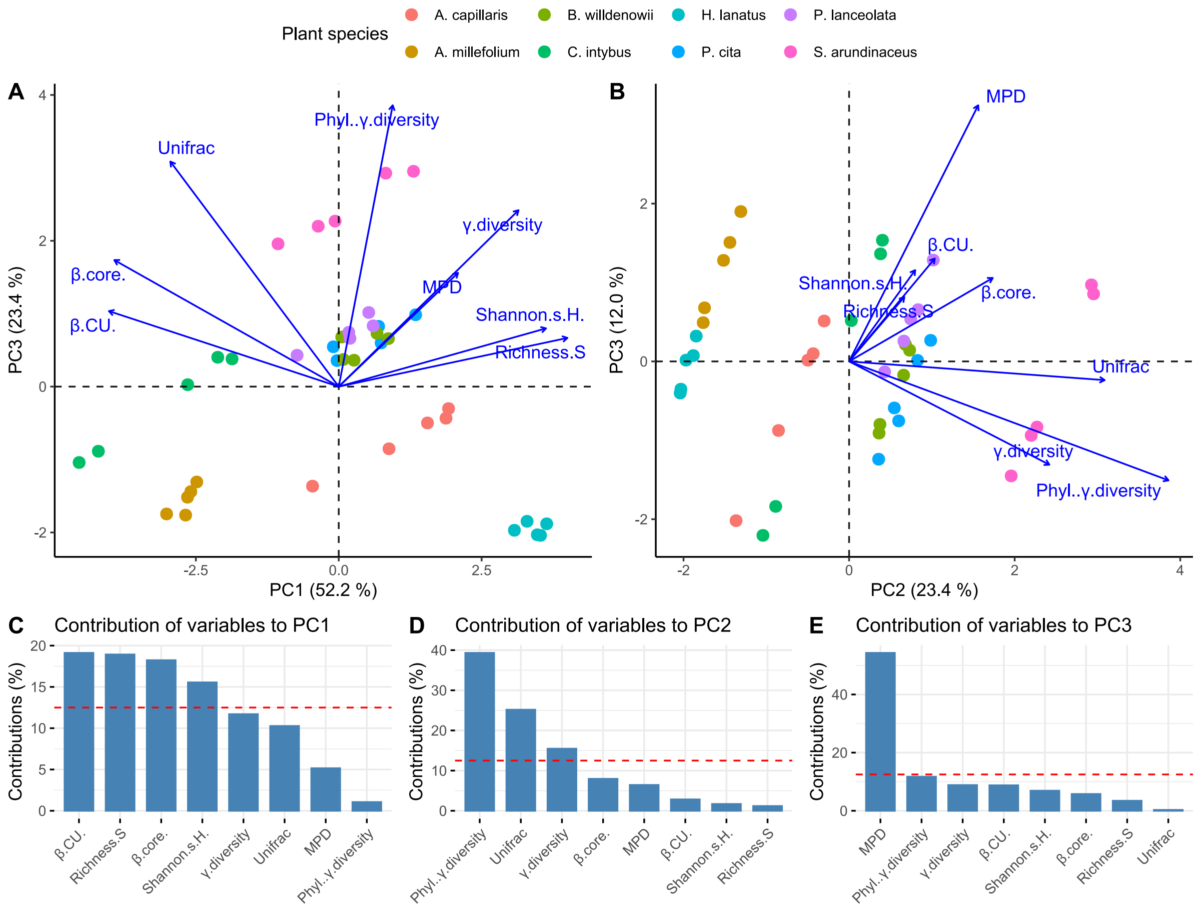

*Table S8: Principal components extracted for all plant individuals. PC = principal component.*

| sample ID | PC1 | PC2 | PC3 | PC4 | PC5 | Plant species |
|---|---|---|---|---|---|---|
| **R01** | -2.48448 | -1.30804 | 1.89947 | -0.78108 | 0.231918 | *A. millefolium* |
| **R02** | -3.01232 | -1.74645 | 0.678831 | -0.16891 | 0.153346 | *A. millefolium* |
| **R05** | -2.67924 | -1.76226 | 0.491428 | 0.913278 | 0.025869 | *A. millefolium* |
| **R09** | -2.58816 | -1.43968 | 1.508185 | -0.26839 | 0.296895 | *A. millefolium* |
| **R10** | -2.64628 | -1.51692 | 1.280865 | -0.18479 | 0.185539 | *A. millefolium* |
| **R12** | -1.86723 | 0.380152 | 1.36305 | 0.675258 | -0.62628 | *C. intybus* |
| **R13** | -4.54268 | -1.04088 | -2.20501 | 0.046957 | -0.34749 | *C. intybus* |
| **R14** | -2.11713 | 0.402471 | 1.535683 | -0.04033 | -0.42621 | *C. intybus* |
| **R15** | -2.64308 | 0.026411 | 0.516775 | 0.376076 | -0.39417 | *C. intybus* |
| **R18** | -4.20632 | -0.8867 | -1.83765 | 0.321026 | -0.3498 | *C. intybus* |
| **R21** | 0.696706 | 0.826869 | 0.012901 | -0.89568 | -0.34062 | *P. cita* |
| **R25** | -0.09488 | 0.546054 | -0.58971 | -1.50472 | 0.009576 | *P. cita* |
| **R26** | 1.344732 | 0.985367 | 0.266758 | 0.344558 | -0.41798 | *P. cita* |
| **R28** | 0.741672 | 0.599315 | -0.75398 | 0.613345 | -0.34203 | *P. cita* |
| **R30** | -0.02944 | 0.357072 | -1.23953 | -0.06613 | 0.064856 | *P. cita* |
| **R34** | -0.06462 | 2.269584 | -0.83228 | 0.504065 | 0.408981 | *S. arundinaceus* |
| **R36** | 0.822038 | 2.927501 | 0.970376 | -0.75853 | 0.079217 | *S. arundinaceus* |
| **R38** | -1.06252 | 1.957826 | -1.45118 | -0.63909 | 0.76841 | *S. arundinaceus* |
| **R39** | 1.307985 | 2.953282 | 0.855307 | 0.683072 | -0.006 | *S. arundinaceus* |
| **R40** | -0.36044 | 2.199818 | -0.93797 | -0.19061 | 0.369722 | *S. arundinaceus* |
| **R42** | 3.637847 | -1.88172 | 0.073857 | -0.02429 | -0.0644 | *H. lanatus* |
| **R44** | 3.073123 | -1.97021 | 0.015929 | -1.40656 | -0.00137 | *H. lanatus* |
| **R45** | 3.530261 | -2.0402 | -0.40109 | 0.433559 | -0.13152 | *H. lanatus* |
| **R46** | 3.294852 | -1.84707 | 0.320811 | -0.81594 | 0.35146 | *H. lanatus* |
| **R47** | 3.47424 | -2.03185 | -0.3524 | 0.322224 | -0.04121 | *H. lanatus* |
| **R51** | 1.921081 | -0.30146 | 0.512232 | 0.572601 | -0.06371 | *A. capillaris* |
| **R53** | -0.45882 | -1.36517 | -2.01851 | -0.12716 | 0.875678 | *A. capillaris* |
| **R54** | 1.552121 | -0.49815 | 0.015444 | 0.649519 | 0.079709 | *A. capillaris* |
| **R55** | 1.876283 | -0.4336 | 0.097731 | 1.105404 | -0.12508 | *A. capillaris* |
| **R59** | 0.879148 | -0.85249 | -0.87613 | 0.759792 | 0.338781 | *A. capillaris* |
| **R61** | 0.668548 | 0.731389 | 0.141992 | -0.68713 | -0.86242 | *B. willdenowii* |
| **R62** | 0.081061 | 0.372582 | -0.79768 | -0.38649 | -0.69525 | *B. willdenowii* |
| **R65** | 0.267182 | 0.361841 | -0.90875 | 0.235152 | -0.76278 | *B. willdenowii* |
| **R66** | 0.86867 | 0.658577 | -0.17467 | 0.061965 | -1.12168 | *B. willdenowii* |
| **R68** | 0.05207 | 0.680063 | 0.215559 | -1.65332 | -0.25871 | *B. willdenowii* |
| **R71** | 0.601928 | 0.833527 | 0.65578 | 1.082188 | 0.408111 | *P. lanceolata* |

| | | | | | | |
|---|---|---|---|---|---|---|
| **R72** | 0.52428 | 1.016759 | 1.284638 | 0.11381 | 0.66873 | *P. lanceolata* |
| **R73** | 0.175394 | 0.74461 | 0.537667 | 0.287495 | 0.5336 | *P. lanceolata* |
| **R77** | 0.195692 | 0.661795 | 0.257931 | 0.784195 | 0.489305 | *P. lanceolata* |
| **R80** | -0.72927 | 0.42999 | -0.13265 | -0.2864 | 1.039011 | *P. lanceolata* |

***Table S9: Variable loadings for the diversity metrics over the first three dimensions.*** *PC = Principal component, β(core) = percentage of AMF present in at least 60% of replicates per plant species, β(CU) = compositional units, MPD = mean phylogenetic diversity.*

| **Diversity metric** | **PC1** | **PC2** | **PC3** |
|---|---|---|---|
| **Richness S** | 0.889 | 0.148 | 0.182 |
| Shannon's H' | 0.806 | 0.178 | 0.257 |
| γ-diversity | 0.699 | 0.538 | -0.291 |
| β(CU) | -0.894 | 0.23 | 0.29 |
| β(core) | -0.873 | 0.386 | 0.235 |
| **UniFrac** | -0.655 | 0.686 | -0.0528 |
| MPD | 0.464 | 0.347 | 0.721 |
| Phylogenetic γ-diversity | 0.21 | 0.858 | -0.335 |

#### Section S5: Bayesian Models

##### AMF biomass

***Figure S5: Posterior Distribution of Bayesian Linear Mixed Model showing the regression coefficients predicting AMF biomass by principal component 3 and root biomass****. The blue bar indicates the mean estimate, and the dark red area indicates the 50% credible interval of each distribution, and the area inside the green lines indicates the 95 % credible interval. See Table S9 for details. Dim.3 = principal component 3 (PC3).*

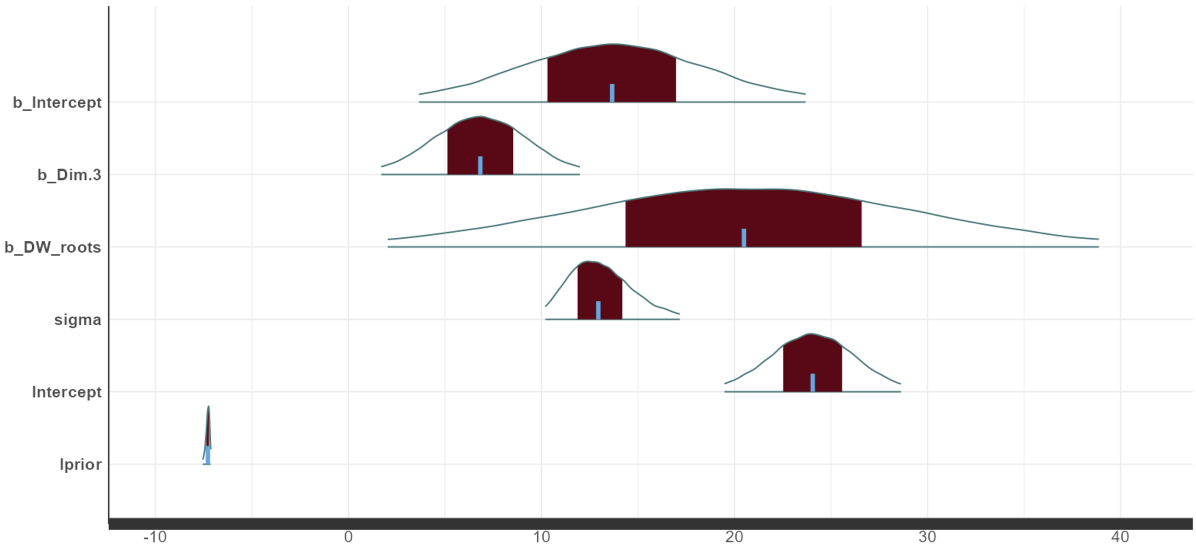

The principal component 3 and the root biomass have a positive relationship with AMF biomass. Especially the distribution of this root biomass as regression coefficient ranges wide (Figure S5, Table S9), indicating that the effect of root biomass on the AMF biomass in the soil is very variable

***Table S10: Point estimates, standard errors and credible intervals of the regression coefficients for the best model after selection: AMF ~ principal component 3 + root biomass****. The model included the plant species identity as random effect. MCSE = Monte Carlo standard error, $\hat{R}$ = $\hat{R}$ −statistic, $e_{eff}$ = effective sample size, CI = Bounds of the central credible interval, representing the central 50(or 97.5)% of the posterior probability distribution, sd = Posterior standard deviation, giving the uncertainty around the estimate mean; σ = Standard deviation of the error.*

| Coefficient | $n_{eff}$ | $\hat{R}$ | mean | MCSE | sd | 2.50% CI | 50% CI | 97.50% CI |
|---|---|---|---|---|---|---|---|---|
| **Intercept** | 56,000 | 1 | 13.66 | 0.02 | 5.07 | 3.65 | 13.66 | 23.69 |
| **PC3** | 56,000 | 1 | 6.83 | 0.011 | 2.59 | 1.7 | 6.83 | 11.99 |
| **root biomass** | 56,000 | 1 | 20.48 | 0.039 | 9.31 | 2.04 | 20.49 | 38.87 |
| **σ** | 45,642 | 1 | 13.14 | 0.008 | 1.79 | 10.2 | 12.95 | 17.17 |
| **Intercept** | 56,000 | 1 | 24.05 | 0.01 | 2.31 | 19.47 | 24.05 | 28.63 |
| **lprior** | 40,247 | 1 | -7.29 | 0.001 | 0.11 | -7.54 | -7.28 | -7.13 |
| **log-posterior** | 23,296 | 1 | -132.08 | 0.01 | 1.52 | -135.93 | -131.73 | -130.2 |

**Table S11: Hypothesis testing of model describing the changes in AMF biomass in the soil.** We test $H_0$: There is evidence to suggest that soil AMF biomass increases with plant interaction generalism. The AMF biomass is measured in nmol per g soil of the neutral lipid fatty acid 16:1ω5. (Alternative: $H_1$: There is no evidence to suggest that biomass increases with plant interaction generalism. The parameter $\beta_{1(2,3)}$ is smaller 0. An evidence ratio of "infinity" implies very large evidence in favor of the hypothesis. Significant model parameters in bold. The evidence ratio is the ratio of the posterior probability of the hypothesis versus its alternative. Models with the random factors were not considered here, as they did not converge. CI = Bounds of the 95% central credible interval.

| *Best fit model:* **AMF biomass ~ $\beta_0$ + $\beta_1$ x PC3 + $\beta_2$ x root DW biomass + σ.** | | | | | | |
|---|---|---|---|---|---|---|
| $H_0$: There is evidence to suggest that biomass increases with plant interaction generalism. The parameter $\beta_1$ is larger than 0, i.e., $\beta_1 > 0$. | | | | | | |
| ***Hypothesis*** | ***Estimate*** | ***Est. Error*** | ***CI Lower*** | ***CI Upper*** | ***Evidence ratio*** | ***Posterior probability*** |
| **$\beta_1$ >0 (PC3)** | **6.83** | **2.59** | **2.59** | **11.09** | **185.05** | **0.99** |
| *Full model:* **AMF biomass ~ $\beta_0$ + $\beta_1$ x PC1 + $\beta_2$ x PC2 + $\beta_3$ x PC3 + $\beta_4$ x root DW biomass + $\beta_5$ x shoot DW biomass σ.** | | | | | | |
| $H_0$: There is evidence to suggest that biomass increases with plant interaction generalism. The parameter $\beta_{1(2,3)}$ is larger than 0, i.e., $\beta_{1(2,3)} > 0$. | | | | | | |
| ***Hypothesis*** | ***Estimate*** | ***Est. Error*** | ***CI Lower*** | ***CI Upper*** | ***Evidence ratio*** | ***Posterior probability*** |
| $\beta_1$ >0 (PC1) | -1.24 | 0.63 | -2.26 | -0.2 | 0.03 | 0.02 |
| $\beta_2$ >0 (PC2) | -0.99 | 0.74 | -2.21 | 0.2 | 0.09 | 0.09 |
| **$\beta_3$ >0 (PC3)** | **7.75** | **1.17** | **5.83** | **9.70** | ∞ | **1** |

***Figure S6:*** *Posterior predictive check comparing the observed outcome variable y to 100 simulated datasets* $y_{rep}$ *from the posterior predictive distribution of the best model predicting the AMF biomass. AMF = Arbuscular mycorrhizal fungi.*

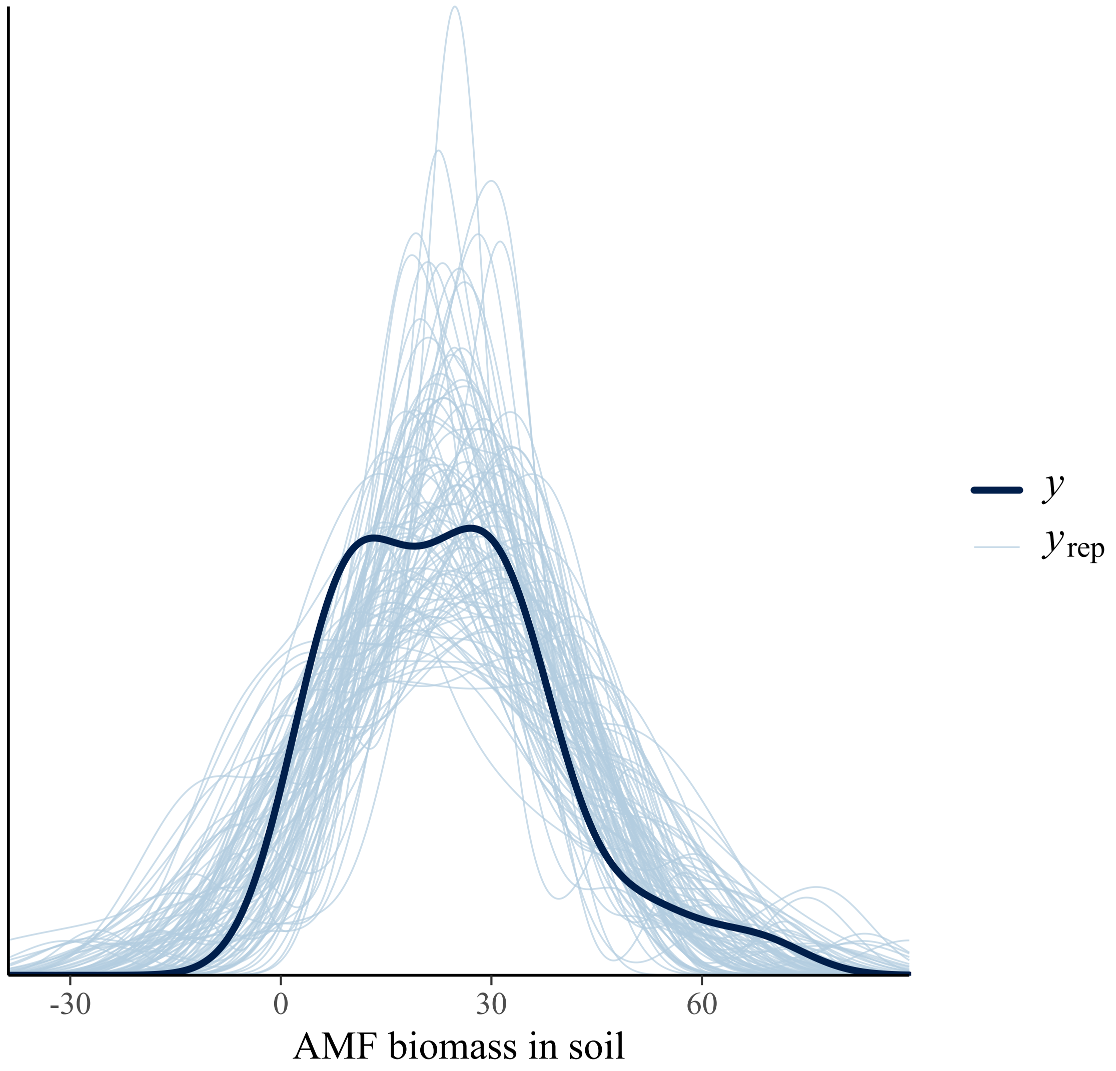

#### Bacterial Biomass

**Figure S7: Posterior Distribution of Bayesian Linear Mixed Model showing the regression coefficients predicting PLFA total bacterial biomass by principal component 1, shoot biomass and root biomass**. *The blue bar indicates the mean estimate, and the dark red area indicates the 50% credible interval of each distribution, and the area inside the green lines indicates the 95 % credible interval. See Table S12 for details. DW_roots = root biomass, Dim,1 = principal component 1, DW_above = shoot biomass, PLFA = Phospholipid fatty acid.*

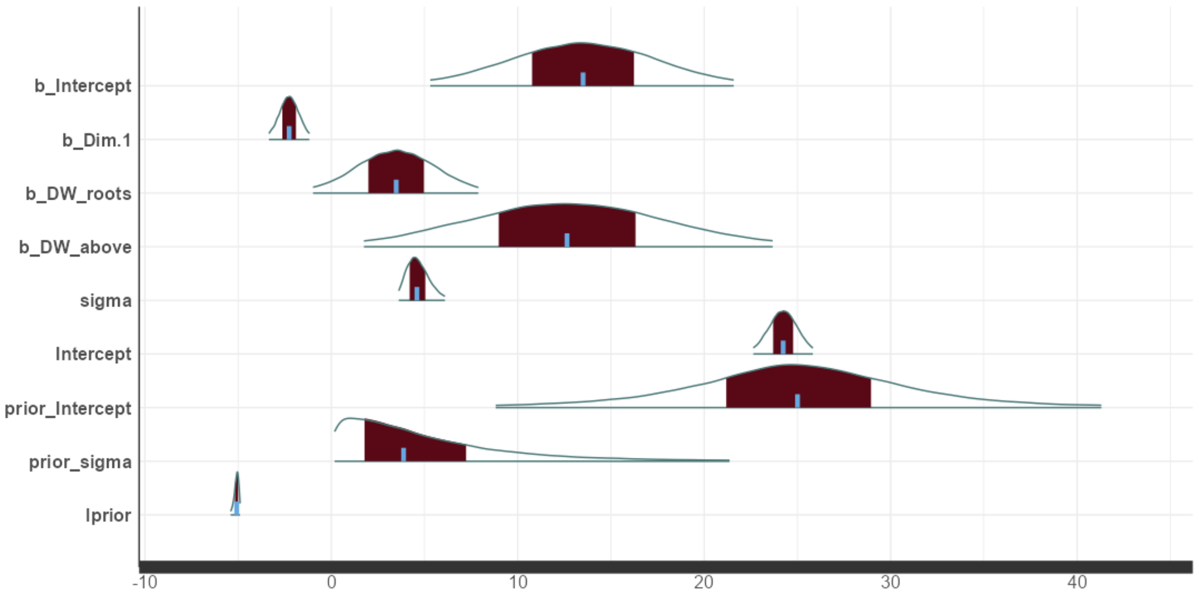

***Table S12: Point estimates, standard errors and credible intervals of the regression coefficients of the best fit Bayesian linear mixed model "soil bacterial biomass ~ PC1 + DW roots + DW shoots".*** *DW = dry weight, MCSE = Monte Carlo standard error,* $\hat{R}$ *=* $\hat{R}$ *−statistic,* $e_{eff}$ *= effective sample size, CI = Bounds of the central credible interval, representing the central 50 (or 97.5)% of the posterior probability distribution, sd = Posterior standard deviation, giving the uncertainty around the estimate mean; σ = Standard deviation of the error.*

| | $n_{eff}$ | $\hat{R}$ | mean | MCSE | sd | 2.50% CI | 50% CI | 97.50% |
|---|---|---|---|---|---|---|---|---|
| **Intercept** | 56,000 | 1 | 13.49 | 0.017 | 4.13 | 5.31 | 13.5 | 21.58 |
| **PC1** | 46,117 | 1 | -2.26 | 0.003 | 0.55 | -3.35 | -2.26 | -1.17 |
| **DW roots** | 56,000 | 1 | 3.47 | 0.009 | 2.25 | -0.98 | 3.48 | 7.89 |
| **DW above** | 48,896 | 1 | 12.65 | 0.025 | 5.55 | 1.75 | 12.64 | 23.67 |
| **σ** | 56,000 | 1 | 4.66 | 0.003 | 0.64 | 3.62 | 4.59 | 6.1 |
| **Intercept** | 56,000 | 1 | 24.24 | 0.003 | 0.81 | 22.64 | 24.24 | 25.83 |
| **prior Intercept** | 56,000 | 1 | 25.01 | 0.038 | 8.99 | 8.82 | 25.01 | 41.3 |
| **prior σ** | 55,242 | 1 | 5.61 | 0.029 | 6.79 | 0.18 | 3.88 | 21.36 |
| **lprior** | 43,308 | 1 | -5.09 | 0.001 | 0.13 | -5.4 | -5.07 | -4.89 |
| **log-posterior** | 20,679 | 1 | -97.89 | 0.012 | 1.7 | -102.11 | -97.55 | -95.64 |

**Table S13: Hypothesis testing of model describing the changes in bacterial biomass in the soil.** We test $H_0$: There is evidence to suggest that soil bacterial biomass increases with plant interaction generalism. The soil bacterial biomass is measured in nmol per g soil of the total bacterial phospholipid fatty acid biomarkers. (Alternative: $H_1$: There is no evidence to suggest that bacterial biomass increases with plant interaction generalism. The parameter $\beta_{1(2,3)}$ is smaller 0). An evidence ratio of "infinity" implies very large evidence in favor of the hypothesis, whereas an evidence ratio of "0" implies very low evidence in favor of the hypothesis. Significant model parameters in bold. Models with the random factors were not considered here, as they did not converge. CI = Bounds of the 95% central credible interval.

| | | | | | | |
|---|---|---|---|---|---|---|
| *Best fit model:* **Bacterial biomass ~ $\beta_0$ + $\beta_1$ x PC1 + $\beta_2$ x root DW biomass + $\beta_5$ x shoot DW biomass + σ.** | | | | | | |
| $H_0$: There is evidence to suggest that bacterial biomass increases with plant interaction generalism. The parameter $\beta_1$ is larger than 0, i.e., $\beta_1 > 0$. | | | | | | |
| ***Hypothesis*** | ***Estimate*** | ***Estimate Error*** | ***CI Lower*** | ***CI Upper*** | ***Evidence ratio*** | ***Posterior probability*** |
| $\beta_1 > 0$ (PC1) | -2.26 | 0.55 | -3.18 | -1.36 | 0 | 0 |
| $\beta_1 < 0$ (PC1) | -2.26 | 0.55 | -3.18 | -1.36 | 18,665.67 | 1 |
| *Full model:* Bacterial **biomass ~ $\beta_0$ + $\beta_1$ x PC1 + $\beta_2$ x PC2 + $\beta_3$ x PC3 + $\beta_4$ x root DW biomass + $\beta_5$ x shoot DW biomass + σ.** | | | | | | |
| $H_0$: There is evidence to suggest that bacterial biomass increases with plant interaction generalism. The parameter $\beta_{1(2,3)}$ is larger than 0, i.e., $\beta_{1(2,3)} > 0$. | | | | | | |
| ***Hypothesis*** | ***Estimate*** | ***Est. Error*** | ***CI Lower*** | ***CI Upper*** | ***Evidence ratio*** | ***Posterior probability*** |
| $\beta_1 > 0$ (PC1) | -2.32 | 0.59 | -3.28 | -1.36 | 0 | 0 |
| $\beta_2 > 0$ (PC2) | -0.04 | 0.71 | -1.20 | 1.12 | 0.9 | 0.47 |

| | | | | | | |
|---|---|---|---|---|---|---|
| $\beta_3$ >0<br>(PC3) | 0.49 | 0.95 | -1.08 | 2.05 | 2.34 | 0.7 |
| ***Alternative:***<br>**$\beta_1$ <0**<br>**(PC1)** | **-2.32** | **0.59** | **-3.28** | **-1.36** | **13,999** | **1** |

***Figure S8: Posterior predictive check comparing the observed outcome variable y to 100 simulated datasets $y_{rep}$ from the posterior predictive distribution of the best model predicting the bacterial biomass.***

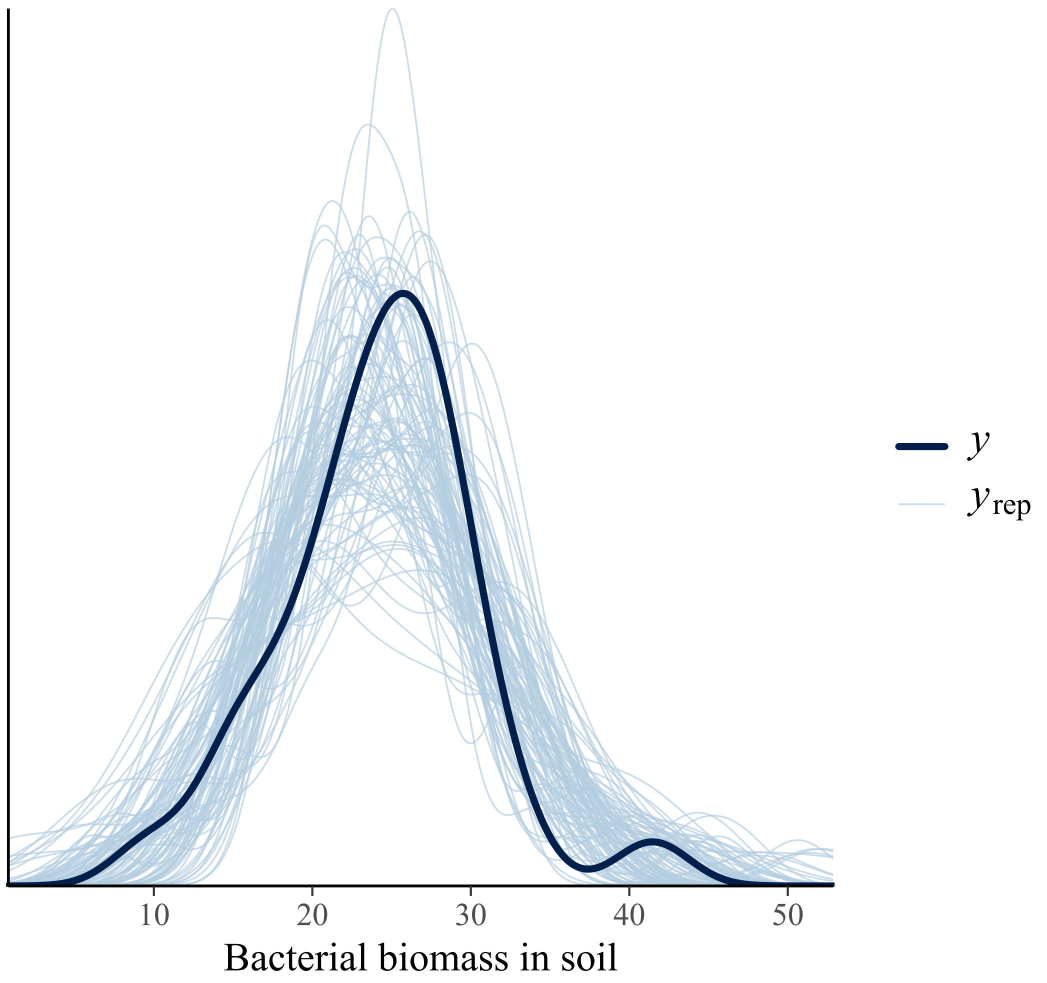

#### Additional models: Mean phylogenetic distance (MPD) as predictor of AMF biomass.

Our best model to predict AMF biomass included PC3, which was strongly informed by MPD. Therefore, we tested if MPD alone is a better predictor than the principal component 3 (PC3) showed that a model that includes both MPD and the root biomass explained with 27% slightly less of the variation than using PC3 as predictor. The difference in predictive accuracy between the two models was tested and showed that these two models were not significantly different from each other.

However, the wide spread of the posterior distribution of MPD as a predictor variable and the inclusion of the value “0” did not indicate that MPD is a better predictor than PC3 of the AMF biomass. Hypothesis testing with $H_0$: “There is evidence to suggest that soil AMF biomass increases with plant interaction generalism.” revealed that MPD significantly increased with soil AMF biomass (evidence ratio = 11,199, posterior probability = 1).

A model that included AMF richness as predictor as well as MPD, root and shoot biomass was computed. While this model that included richness explained with 30% slightly more of the variance than when excluding richness, a model comparison using the leave-one-out algorithm did not show a significant difference between the models. Furthermore, hypothesis testing revealed that there was no evidence that the model parameter associated with richness differed from “0”.

Models that included richness but not MPD explained less of the variance of the AMF biomass and hypothesis testing if richness showed that richness was not a significant predictor in these models.

**Figure S9: Posterior Distribution of Bayesian Linear Mixed Model showing the regression coefficients predicting soil AMF biomass by mean phylogenetic distance (MPD) and root biomass**. *The blue bar indicates the mean estimate, and the dark red area indicates the 50% credible interval of each distribution, and the area inside the green lines indicates the 95 % credible interval. See Table S12 for details. DW_roots = root biomass, AMF = Arbuscular mycorrhizal fungi.*

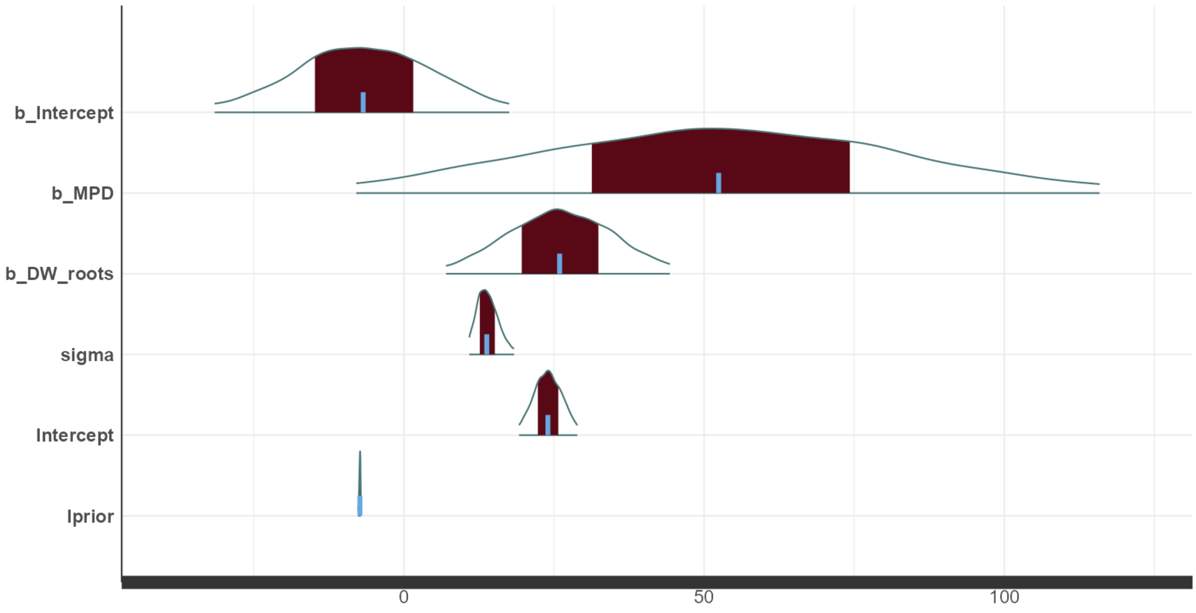

***Table S14: Point estimates, standard errors and credible intervals of the regression coefficients of the best fit Bayesian linear mixed model "soil AMF biomass ~ MPD + DW roots".*** *DW = dry weight, MCSE = Monte Carlo standard error,* $\hat{R} = \hat{R}$ *−statistic,* $e_{eff}$ *= effective sample size, CI = Bounds of the central credible interval, representing the central 50 (and 97.5)% of the posterior probability distribution, sd = Posterior standard deviation, giving the uncertainty around the estimate mean; σ = Standard deviation of the error.*

| | $n_{eff}$ | $\hat{R}$ | mean | MCSE | sd | 2.50% CI | 50% CI | 97.50% |
|---|---|---|---|---|---|---|---|---|
| b_Intercept | 56,000 | 1 | -6.04 | 0.023 | 5.42 | -16.63 | -6.07 | 4.62 |
| b_MPD | 56,000 | 1 | 55.46 | 0.059 | 13.86 | 28.23 | 55.51 | 82.48 |
| b_DW_roots | 56,000 | 1 | 24.38 | 0.018 | 4.26 | 16.02 | 24.38 | 32.76 |
| sigma | 56,000 | 1 | 13.25 | 0.003 | 0.78 | 11.83 | 13.22 | 14.89 |
| Intercept | 56,000 | 1 | 25.23 | 0.005 | 1.09 | 23.09 | 25.24 | 27.39 |
| lprior | 56,000 | 1 | -7.21 | 0 | 0.05 | -7.31 | -7.2 | -7.12 |
| log-posterior | 28,633 | 1 | -592.74 | 0.009 | 1.44 | -596.42 | -592.41 | -590.95 |

***Figure S10:*** *Predicted changes AMF biomass (NLFA 16:1ω5) in rhizosphere soil as a function of mean phylogenetic distance (MPD) and root biomass (n=32). Root biomass was modelled as covariate, but here is visualized at three levels (mean root biomass ± standard deviation, uncertainty interval = 0.5). Final model: AMF biomass ~ MPD + root biomass.*

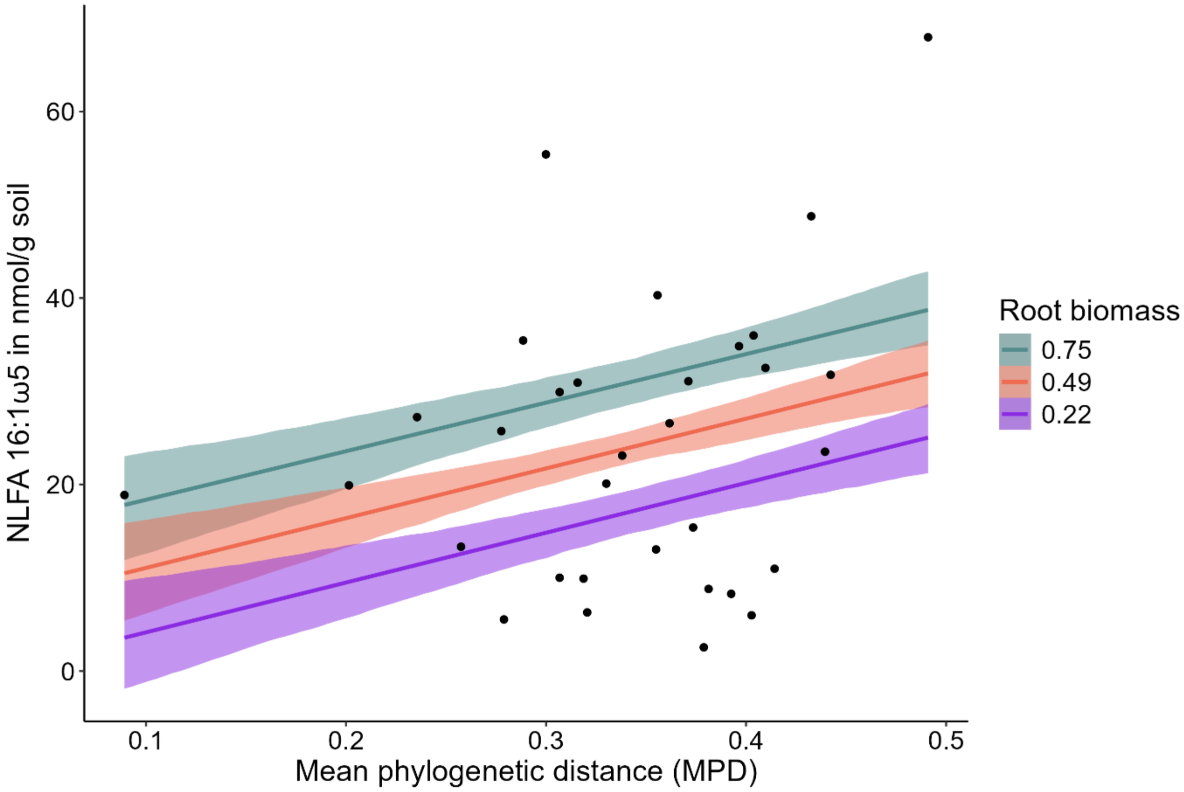

***Figure S11: Posterior predictive check comparing the observed outcome variable y to 100 simulated datasets $y_{rep}$ from the posterior predictive distribution of the best model predicting the bacterial biomass.***

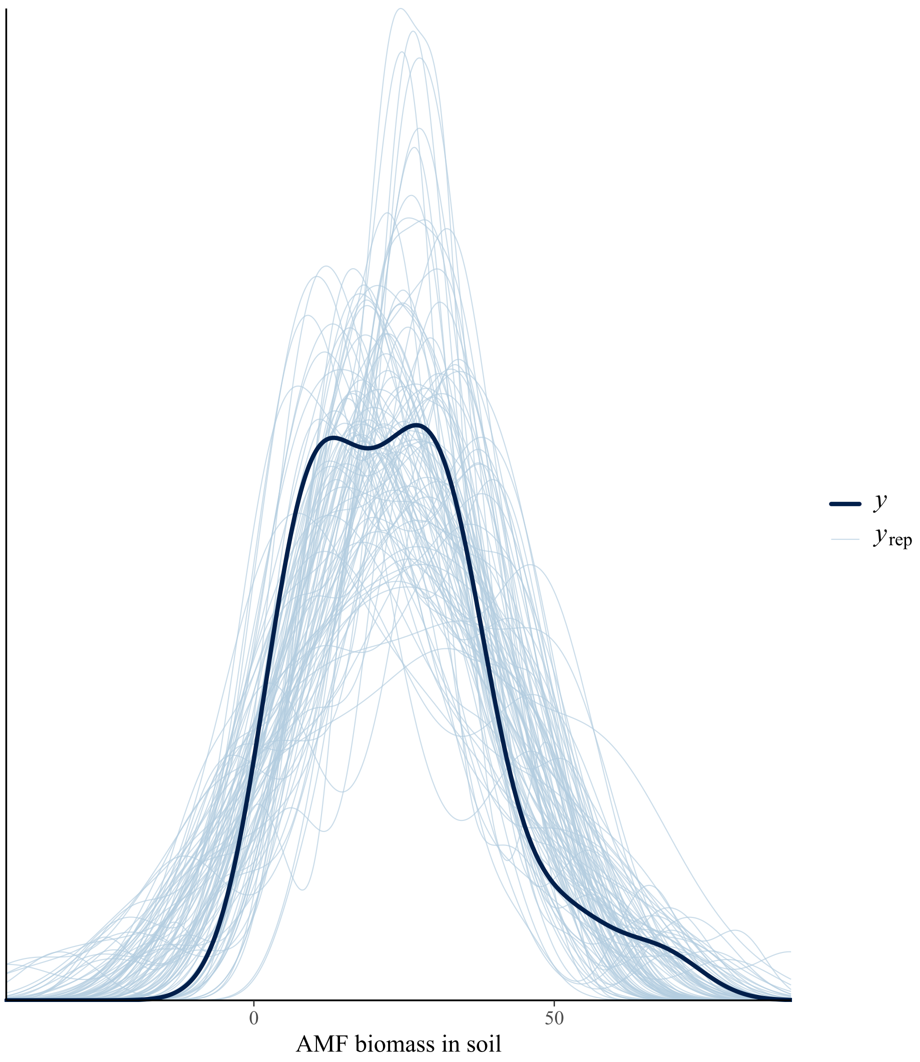

#### Additional models: AMF richness as predictor of total bacterial biomass in the soil

The richness of the AMF interacting with each plant had a strong impact on principal component 1 which predicted the soil bacterial biomass best. Testing if richness alone is a better predictor for soil bacterial biomass than PC1, overall indicated that PC1 was a better predicter. Richness, and both the root and shoot biomass explained with 32% less of the variance than using PC1 as predictor instead of richness. Testing the difference in predictive accuracy between the two models with the leave one out (loo) function revealed significant differences, with the model using PC1 as predictor showing higher accuracy. We also tested the hypothesis $H_0$: "There is evidence to suggest that soil bacterial biomass increases with AMF richness.", which revealed that bacterial biomass in the soil did not increase with richness. Instead, richness had a significant negative model parameter (evidence ratio = 64.57, posterior probability = 0.98). For details see Figures S12 –

**Figure S12: Posterior Distribution of Bayesian Linear Model showing the regression coefficients predicting soil bacterial biomass by richness S, root and shoot biomass**. *The blue bar indicates the mean estimate, and the dark red area indicates the 50% credible interval of each distribution, and the area inside the green lines indicates the 95 % credible interval. See Table S  for details. DW_roots = root biomass, DW_above = shoot biomass.*

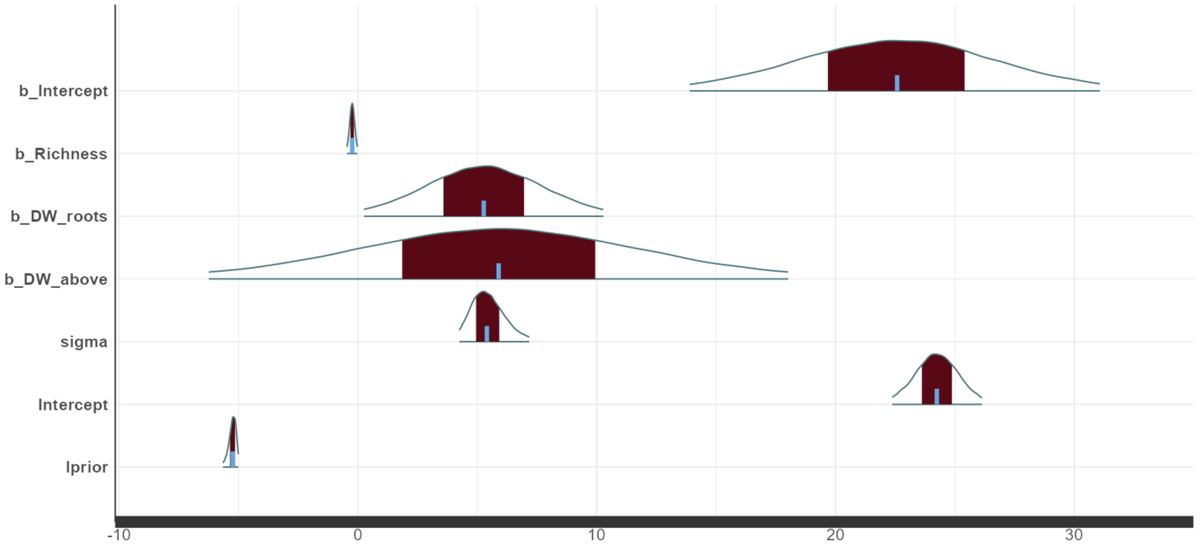

***Table S15: Point estimates, standard errors and credible intervals of the regression coefficients of the best fit Bayesian linear model "soil bacterial biomass ~ Richness + root biomass + shoot biomass".*** *DW = dry weight, MCSE = Monte Carlo standard error,* $\hat{R}$ *=* $\hat{R}$ *−statistic,* $e_{eff}$ *= effective sample size, CI = Bounds of the central credible interval, representing the central 50 (and 97.5)% of the posterior probability distribution, sd = Posterior standard deviation, giving the uncertainty around the estimate mean; σ = Standard deviation of the error.*

| | $n_{eff}$ | $\hat{R}$ | mean | MCSE | sd | 2.50% CI | 50% CI | 97.50% |
|---|---|---|---|---|---|---|---|---|
| **Intercept** | 56,000 | 1 | 22.55 | 0.018 | 4.35 | 13.89 | 22.58 | 31.07 |
| **Richness** | 52,506 | 1 | -0.24 | 0 | 0.11 | -0.45 | -0.24 | -0.02 |
| **DW_roots** | 56,000 | 1 | 5.27 | 0.011 | 2.55 | 0.26 | 5.27 | 10.28 |
| **DW_above** | 56,000 | 1 | 5.9 | 0.026 | 6.14 | -6.24 | 5.9 | 18.02 |
| **σ** | 56,000 | 1 | 5.48 | 0.003 | 0.75 | 4.25 | 5.4 | 7.18 |
| **Prior Intercept** | 56,000 | 1 | 24.24 | 0.004 | 0.95 | 22.37 | 24.24 | 26.13 |
| **lprior** | 42,782 | 1 | -5.26 | 0.001 | 0.17 | -5.65 | -5.24 | -5 |
| **log-posterior** | 22,892 | 1 | -103.18 | 0.011 | 1.71 | -107.39 | -102.84 | -100.93 |

*Figure S10: Predicted changes in soil bacterial biomass as a function of AMMF richness modelled for three levels of shoot and root biomass, uncertainty interval = 0.5 (n = 32). Best model: soil bacterial biomass ~ richness + root biomass + shoot biomass.*

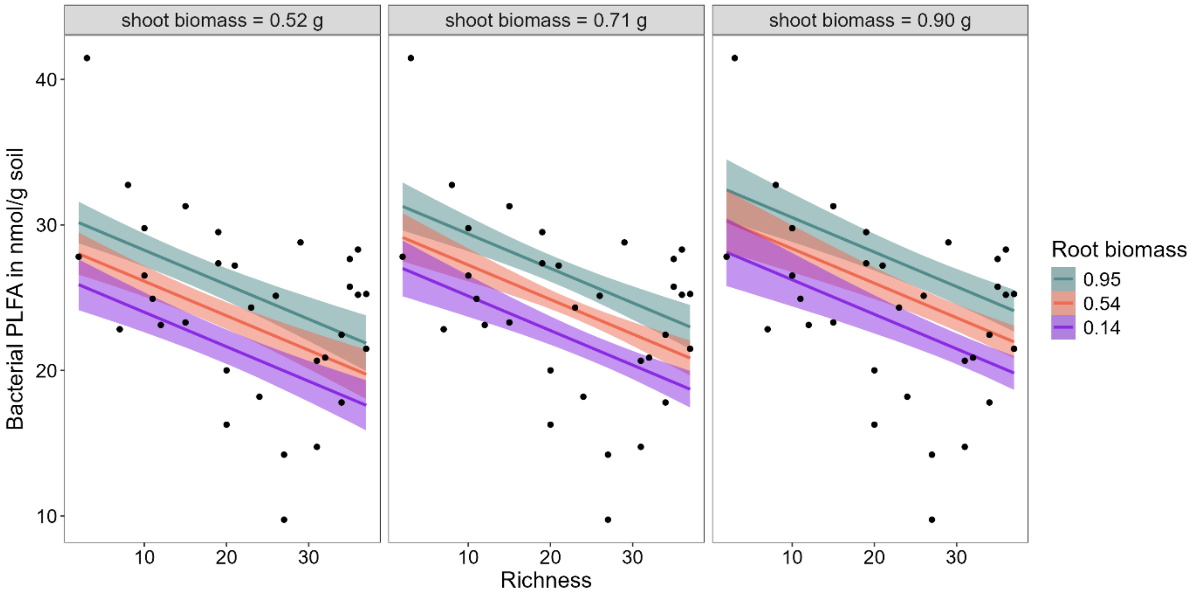

***Figure S: Posterior predictive check comparing the observed outcome variable y to 100 simulated datasets $y_{rep}$ from the posterior predictive distribution of the best model predicting the bacterial biomass.***

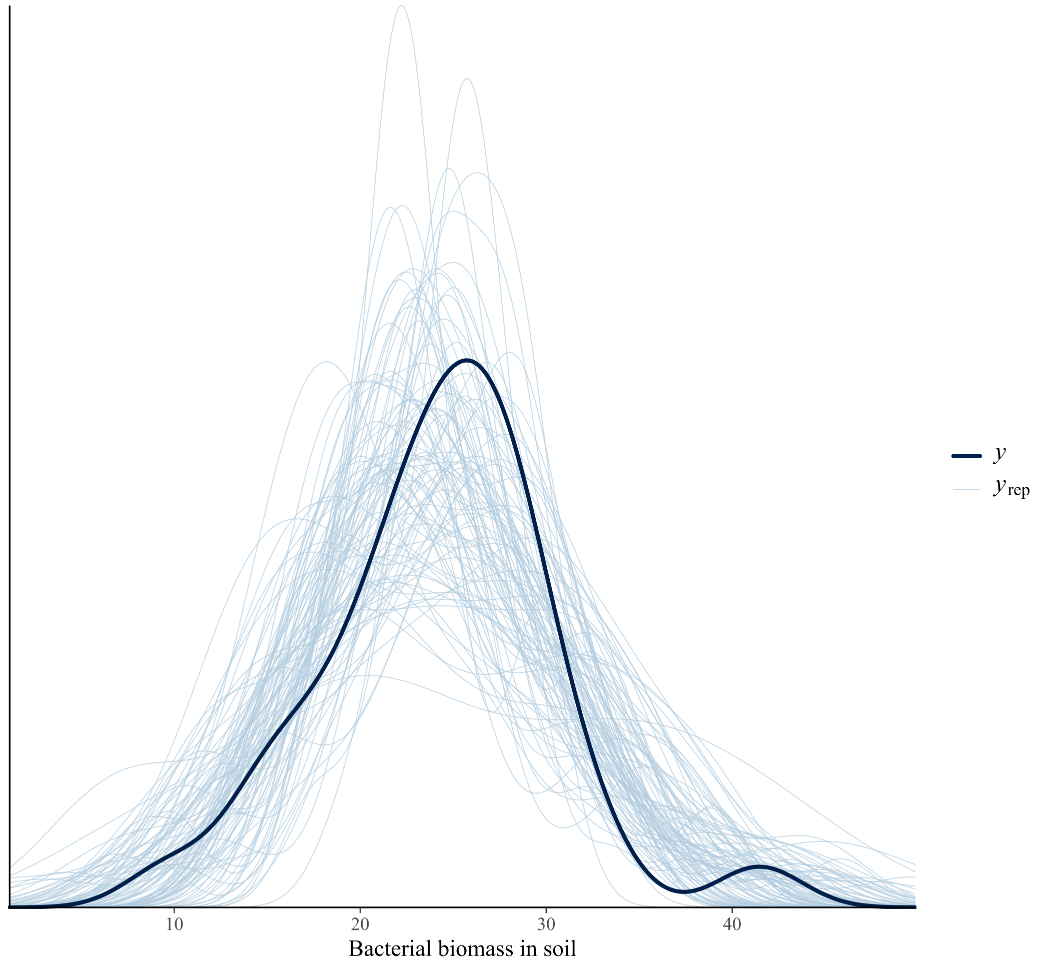
